# Apparent thinning of visual cortex during childhood is associated with myelination, not pruning

**DOI:** 10.1101/368274

**Authors:** Vaidehi S. Natu, Jesse Gomez, Michael Barnett, Brianna Jeska, Evgeniya Kirilina, Carsten Jaeger, Zonglei Zhen, Siobhan Cox, Kevin S. Weiner, Nikolaus Weiskopf, Kalanit Grill-Spector

## Abstract

Microstructural mechanisms underlying apparent cortical thinning during childhood development are unknown. Using functional, quantitative, and diffusion magnetic resonance imaging in children and adults, we tested if tissue growth (lower T1 relaxation time and mean diffusivity (MD)) or pruning (higher T1 and MD) underlies cortical thinning in ventral temporal cortex (VTC). After age 5, T_1_ and MD decreased in mid and deep cortex of functionally-defined regions in lateral VTC, and in their adjacent white matter. T1 and MD decreases were (i) consistent with tissue growth related to myelin proliferation, which we verified with adult postmortem histology and (ii) correlated with apparent cortical thinning. Thus, contrary to prevailing theories, cortical tissue does not thin during childhood, it becomes more myelinated, shifting the gray-white matter boundary deeper into cortex. As tissue growth is prominent in regions with protracted functional development, our data suggest an intriguing hypothesis that functional development and myelination are interlinked.

---

As the brain develops from infancy to adulthood, magnetic resonance imaging (MRI) shows that cortex thins^1-6^. Early sensory regions thin before higher-level frontal and temporal regions^1,3^. However, the mechanisms underlying cortical thinning during development are not well understood.

Three developmental theories have been proposed to explain cortical thinning across development: pruning, myelination, and cortical morphology. *Pruning*, evaluated using cross-sectional histological studies on post mortem brains, suggests that the removal of inefficient synapses, dendrites, and neurons leads to cortical tissue loss^7-9^. This subsequently produces thinner cortex in adulthood. Pruning is thought to improve neural processing by optimizing brain circuits for particular operations. *Myelination* suggests that the myelin sheath wrapping axons grows during development^2, 10-13^, thus increasing the efficiency of saltatory conduction, and leading to faster and more reliable information transmission. As myelin increases raise the intensity of voxels in T_1_-weighted anatomical images, voxels close to the gray-white matter boundary will appear brighter in adults than children, thus shifting the gray-white boundary deeper into cortex in adults^14^. *Cortical morphology* (cortical folding and surface area) suggests that during development sulcification and surface area increase due to mechanical forces^15,16^, thereby producing thinner cortex in adulthood^17^. These mechanisms are not mutually exclusive as a combination of pruning, increased myelination, and morphological alterations may result in thinner cortex in adulthood.

Advances in quantitative MRI (qMRI)^18-20^ and diffusion MRI (dMRI)^21^ provide independent and complementary non-invasive, measurements of gray and white matter tissue properties, allowing us to disambiguate developmental hypotheses in the living human brain. Quantitative MRI enables the measurement and comparison across individuals of the amount of non-water tissue within a voxel (macromolecular tissue volume, MTV) and the relaxation time (T_1_), which depend on tissue composition (e.g., tissue containing myelin, reduces T_1_ more than tissue without it). Additionally, mean diffusivity (MD), obtained from dMRI, depends on the size, density, and structure of the space within tissue through which water diffuses and provides additional insight into microstructural changes during development^20,22^.

*How can these MRI measurements differentiate the three developmental hypotheses?* Although we cannot measure microstructure directly using *in vivo* MRI, we can distinguish these theories because their effects on microstructure within a voxel differs. Pruning results in developmental reductions in synaptic spines^7^, dendrites, and neurons. Although, the magnitude of the effect of pruning on T_1_ or MD remains unknown, pruning results in less cortical tissue for protons to exchange energy with and less hindrance on diffusion of water molecules, predicting higher T_1_ and MD in adult’s cortex than children’s. In contrast, myelination predicts developmental changes to both white and gray matter. In the white matter, increased myelination predicts lower T_1_^18,23^ and reduced MD^20,22^. Myelination is most pronounced in deep cortical layers closer to gray-white boundary, which contains more afferent and efferent projections, than layers close to pial surface^24, 25^. Thus, in gray matter, developmental myelination predicts lower T_1_^26,27^ and lower MD in adult’s than children’s cortex, especially in deeper layers. Finally, developmental changes in cortical morphology predict no development of either T_1_ or MD of gray or white matter. Instead it predicts morphological changes in the local cortical curvature and surface area.

We applied these novel methods to address developmental theories of cortical thinning. We obtained multiple independent measures of fMRI, qMRI, and dMRI in 27 children (ages 5-9, *N*=12, 9 female; ages 10-12, *N*=15, 5 female) and 30 adults (ages 22-28, 11 female). We used fMRI to define functional regions of interest (fROIs) selective to faces, characters, and places within ventral temporal cortex (VTC) of each participant using a functional localizer experiment (**Fig. S1a**). We focused on these VTC regions for two reasons. First, accumulating evidence shows differential functional^28-31^ and anatomical development^26^ across VTC, whereby face- and character-selective regions show prolonged development compared to place-selective regions. Thus, examination of the development of cortical thickness (CT) within VTC allows testing if thinning is guided by uniform or heterogeneous mechanisms across a cortical expanse. Second, functional regions within VTC can be identified in each participant’s brain without spatial smoothing, and have a reliable arrangement with respect to the cortical folding^32, 33^. Thus, studying VTC offers enhanced precision for examining developmental mechanisms of thinning and an opportunity to test the relationship between cortical thinning and morphology. After localizing functional regions of interest (fROIs) in each participant, we measured: (1) CT of each fROI, (2) T_1_ and MD in white matter adjacent to each fROI using qMRI and dMRI, respectively, (3) T_1_ and MD in gray matter of each fROI across intra-cortical depths, and (4) cortical curvature of each fROI. We compared these measurements across age groups to determine which factors develop and if so, whether these developments are region-specific or region-general. Finally, we tested if cortical thinning is linked to development of T_1_, MD, or cortical curvature.

## Results

First, we verified that data quality was not lower in children than adults. Due to excessive motion (> 2 voxels) during fMRI, data from 3 out of 30 adults and 1 out of 27 children were removed from further analysis. In the remaining 26 children and 27 adults, we found no significant differences in motion during scan (F2,100=2.61, *p*>0.05, **Fig. S1b**). fMRI timeseries signal-to-noise ratio (tSNR) across VTC was higher in children than adults (main effect of age: F2,100=37.49, *p*<0.05, **Fig. S1c**). For dMRI, data from 2 children, 4 preteens, and 3 adults were excluded because of excessive motion above 2 voxels (1 voxel = 2 × 2 × 2 mm^3^), leaving 20 children (ages 5-9, *N=*9, ages 10-12, *N*=11) and 24 adults (ages 22-28) for dMRI analyses. After these exclusions, there was no significant difference across age groups. These quality control analyses demonstrate that (i) we can obtain high quality measurements in children and (ii) any developments that we report are likely not driven by nonspecific differences between age groups such as head motion or lower tSNR.

In each participant’s brain, we defined fROIs selective for faces (pFus-faces and mFus-faces, which together are referred to as the fusiform face area^34,35^), characters (pOTS-chars and mOTS-chars located in the occipital temporal sulcus (OTS), and also referred as visual word form area, VWFA, 1 and 2, respectively^36, 37^), and places (CoS-places, also referred as parahippocampal place area^38^) from the localizer experiment. We were able to localize these fROIs even in our youngest participants (**Fig. 1a** and **Supplemental Table 1**). Additionally, in each subject we generated whole brain maps of T_1_ and MD, and measured mean T_1_ and MD values in each subject’s fROIs and their adjacent white matter.

**Figure 1.**
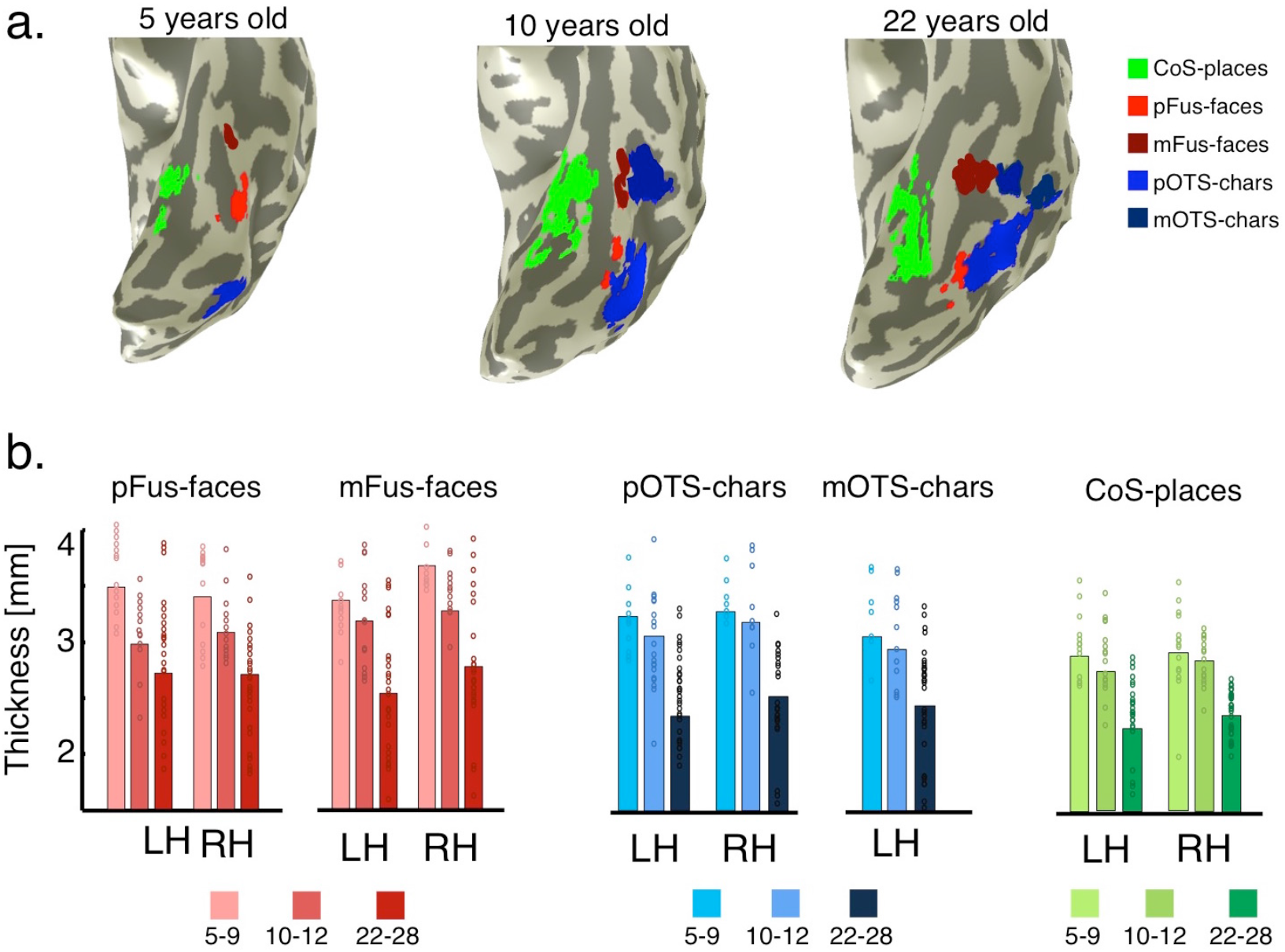
Cortical thickness (CT) decreases from age 5 to adulthood in category-selective regions of ventral temporal cortex (VTC). (a) Functional regions in the left ventral temporal cortex (VTC) of three example subjects, ages 5, 10, and 22 years. fROIs were defined by contrasting responses to one category versus others, with a t>3, voxel-level threshold. (b) Average CT in face-selective regions in the lateral fusiform gyrus (pFus-faces and mFus-faces, which together are also referred to as the fusiform face area, FFA), character-selective regions in the occipito-temporal sulcus, OTS, (pOTS-characters and mOTS- characters, which are also referred to as the visual word form area, VWFA), and a place-selective region in the collateral sulcus (CoS-places, also referred to as the parahippocampal place area, PPA) across children (ages 5-9, *N*=11, 10-12, *N*=15) and adults (ages 22-28 years, N=27). CT is significantly higher in children than adults in all fROIs (F2,384=78.74, *p*<0.05). *Bar height:* average CT; *Dots:* individual subject’s CT. *LH:* left hemisphere. *RH:* right hemisphere.

### VTC thins from childhood to adulthood

Maps of CT on each subject’s cortical surface were generated using FreeSurfer^39^ from T_1_ images. Our measurements of CT are consistent with prior *ex vivo*^40^ and *in vivo* studies^2^ (**Fig. S2**). In general, CT decreased with age across VTC, except for a region near the temporal pole in which CT increased with age (interaction between age of subject and anatomical parcel, *F*16,900 = 19.48, *p*<0.05, **Figs. S3a,b**).

Measurements of CT in fROIs of VTC showed that cortex thinned from age 5 to adulthood in face-, character-, and place-selective regions (main effect of age, 3-way repeated measures analysis of variance (ANOVA) with factors: age of subject (5-9/10-12/22-28 years), fROI (face-/place-/character-selective), and hemisphere (left/right), F2,384 =78.74, *p*<0.05, **Fig. 1b**). Children’s cortex (ages 5-9) was on average 0.73±0.11 mm thicker than adults’. The smallest difference was observed in right CoS-places (0.46 mm) and the largest difference in right mFus-faces (0.97 mm). Additionally, CT varied across fROIs (main effect of fROI, F2,384 =21.00, *p*<0.05), whereby cortex was thinnest in CoS-places (2.56±0.45 mm), which lies close to the fundus of the collateral sulcus (CoS), and thickest in mFus-faces (2.97±0.65mm), which lies on the lateral fusiform gyrus. There was also a small but significant hemispheric difference, as left hemisphere fROIs were thinner than the right ones (F1,384 =4.99, *p*<0.05). As cortex thins, the volume of all fROIs did not decrease. There was no significant development in the volume of character- and place-selective regions (*p=n.s*.), and a developmental increase in the volume of face-selective regions (F2,206=5.2, *p*<0.05).

### In functionally defined white matter (FDWM), T_1_ relaxation time and MD decrease from childhood to adulthood

As we found substantial development in CT, we first examined if these developments may be related to changes in white matter properties adjacent to each fROI. Therefore, we dilated each fROI 5 mm into the white matter (**Fig. 2a** and Methods), which we refer to as functionally-defined-white-matter (FDWM^41^). We estimated T1 (**Fig.2b**) and MD (**Fig. 2c**) in FDWM of face-, character-, and place-selective fROIs in each participant. Results were similar for fROIs from the same domain. Thus, in subsequent main figures we show a single fROI from each domain: face-selective mFus-faces, character-selective mOTS- chars, and place-selective CoS-places. Data from additional face (pFus-faces) and character fROIs (pOTS- chars) are in **Figs. S4, S6**.

Results show differential development of T1 in FDWM (interaction between age of subject and fROI, F4,384=2.54, *p*<0.05, **Figs. 2d** and **S4a**). T_1_ progressively decreased from age 5-9 to 10-12 to adulthood in the FDWM near face- and character-selective regions (Fs>18.12, *p*s<0.05). However, there was no significant change in T_1_ in FDWM of place-selective cortex after age 5 (F_2,98_=2.3, *p*=n.s.) In FDWM where T1 decreased, MTV concomitantly significantly increased with age. (**Fig. S5**).

**Figure 2.**
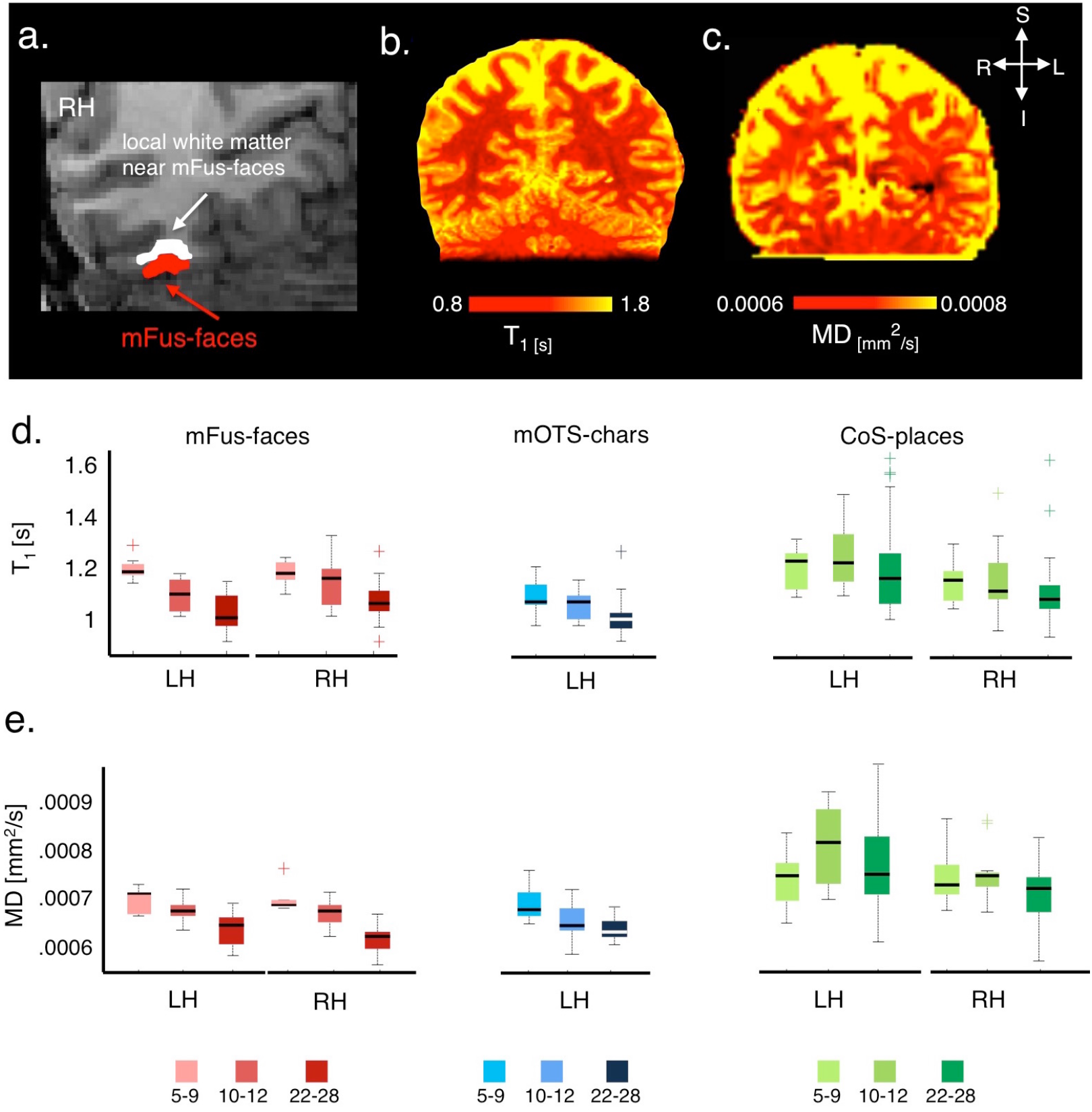
Development of T_1_ relaxation time and mean diffusivity (MD) in functionally-defined white matter (FDWM) of VTC. (a) Section of a coronal slice showing face-selective mFus-faces (red) in the right hemisphere of an example adult brain and its associated FDWM (white), which was generated by dilating the fROI into the adjacent white matter. (b-c) Coronal slice in an example adult brain showing T_1_ and MD maps, respectively. T_1_ and MD are lower in myelinated white matter than in the gray matter. (d) Box plots showing median (thick line), 25^th^ and 75^th^ percentile (box), and range (whiskers) of T_1_ across participants of each age group in bilateral mFus-faces (red), left mOTS-characters (blue), and bilateral CoS-places (green). Crosshairs indicate outliers. T_1_ progressively decreased from ages 5-9 (*N*=11) to 10-12 (*N*=15) to adulthood (*N*=27) in face- and character-selective regions, but not in CoS-places. (e) Box plots showing MD in the same fROIs. Like T_1_, MD also progressively decreased from ages 5-9 (*N*=9) to 10-12 (*N*=11) to adulthood (*N*=24) in face- and character-selective regions, but not in CoS-places. In all box plots: *light colors:*5-9 year olds; *medium colors:* 10-12 year olds, and *dark colors:* 22-28 year olds. *LH:* left hemisphere. *RH:* right hemisphere.

MD in FDWM also revealed differential development (interaction between age of subject and fROI, F4,320=4.64, *p*<0.05). MD monotonically decreased in FDWM near face- and character-selective fROIs from age 5 to adulthood (Fs>8.77, *p*s<0.05, **Figs. 2e** and **S4b**). However, there was no development in MD of FDWM near CoS-places (F_2,80_=2.57, *p*>0.05). Decreases in both T_1_ and MD in FDWM next to face- and character-selective fROIs (but not place-selective cortex) are consistent with the hypothesis that development of white matter near these fROIs is associated with increased myelination.

### In mid and deep cortical depths, T_1_ and MD decrease from childhood to adulthood

Next, we tested if there are tissue developments in cortex. We evaluated T_1_ and MD in each fROI across cortical depths in equidistant steps (10 steps for T_1_ and 8 steps for MD), extending from the pial surface to gray-white matter boundary, with the last two steps inside the white matter (**Fig. S6**).

Data reveal three findings. First, mean T_1_ decreased with age in face- and character-selective cortex (Fs>4.37, *p*<0.05), but not in place-selective cortex (F_2,98_=0.35, *p*=n.s.). Second, examination of T_1_ across cortical depth shows that T_1_ near the gray-white matter boundary was lower than T_1_ in superficial layers of gray matter (main effect of cortical depth in a 4-way ANOVA with factors of age of subject, fROI, hemisphere, and depth, **see online methods**, F4,1920 = 3450.47, *p*<0.05, **Fig. 3a**). Third, across cortical depths, in face- and character-selective fROIs, the largest development in T_1_ occurred away from the superficial pial surface and was more prominent in mid cortical depths (interaction between age of subject and depth in 4-way ANOVA with factors of age of subject, fROI, hemisphere, and depth, F8,1920 = 17.51, *p*<0.05, **Fig. 3a**). In contrast, in CoS-places, T_1_ curves largely overlapped across age groups. Data from pFus-faces and pOTS-chars are in **Fig. S7a**.

Notably, T_1_ curves were shifted leftward in adults compared to children. In other words, comparable T1 values were more superficial in adults than children. For example, T_1_ at the gray-white boundary in 5–9 year-olds was equivalent to T_1_ at ∼80% depth in adults. To quantify this shift, we fitted a maximum log-likelihood function to each subject’s T_1_ curve, and estimated the inflection point of the curve, which reflects the depth at which slope is maximal (**Fig. 3b**). Results revealed that the depth of this inflection point was closer to the pial surface in adults than children, with larger development in face- and character-selective fROIs than in place-selective fROIs (significant interaction between age of subject and fROI, F4,384 = 3.89, *p*<0.05).

**Figure 3.**
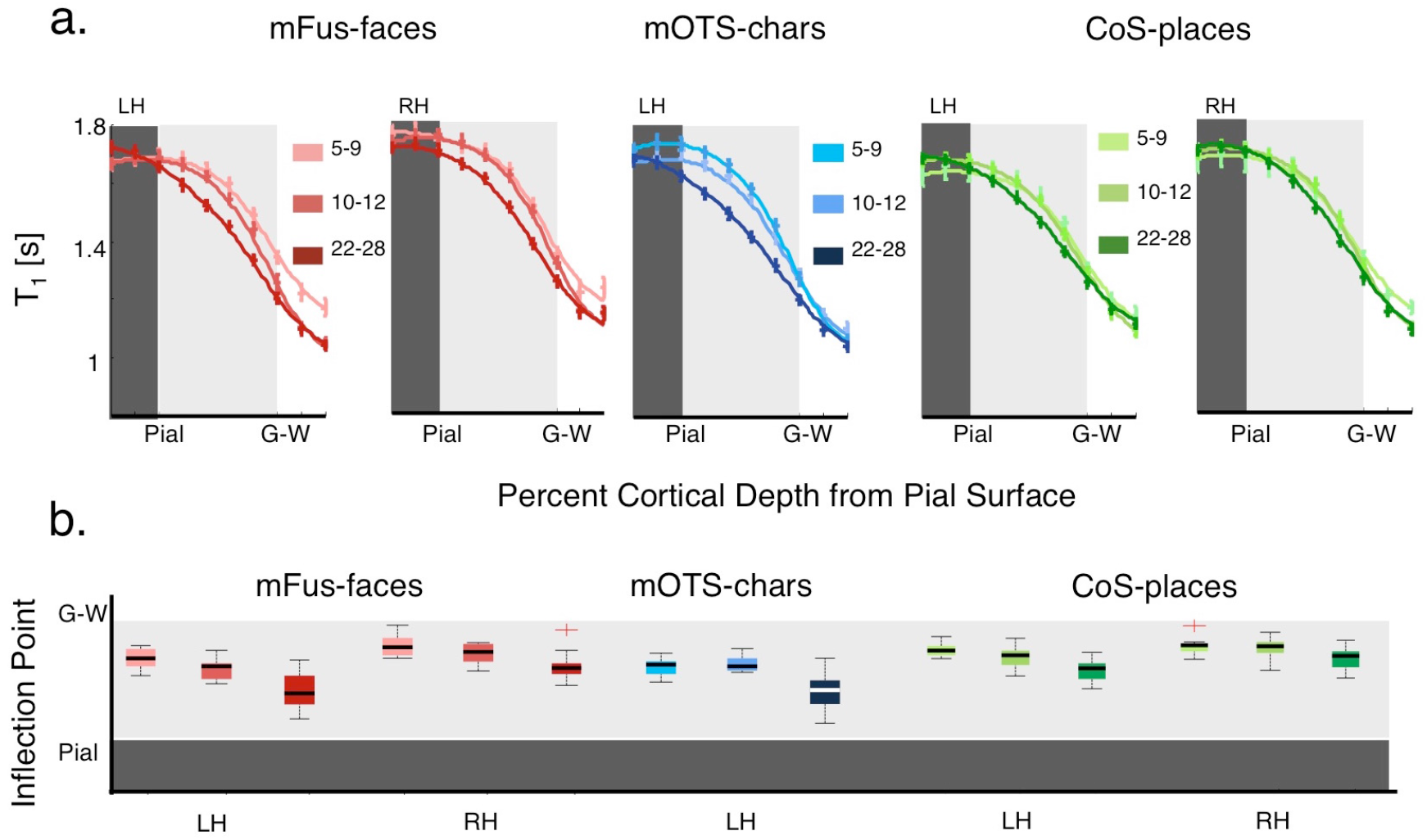
Development of T_1_ in gray matter as a function of cortical depth. (a) T1 curves across equidistant intra-cortical depths from the pial surface into adjacent white matter tissue in bilateral mFus- faces (red), left mOTS-characters (blue), and bilateral CoS-places (green) across the three age groups. (Same subjects as **Fig. 2d**). *Errorbars:* standard error of the mean (SEM) across participants within an age group. (b) Box plots of the inflection point of T_1_ curves across age groups. The inflection point reflects the cortical depth at which slope of the T_1_ curve is maximal. *LH:* left hemisphere. *RH:* right hemisphere. Same conventions as **Fig 2**.

The development of MD across intra-cortical depths is more complex as gray matter is not as directionally organized as white matter (**Figs. S7b, S8a**). Across age groups, MD decreased from the pial surface to the gray-white matter boundary (interaction between age of subject and cortical depth: F6,1248=8.26, *p*<0.05). Analysis of MD revealed differential development across fROIs (significant interaction between age of subject and fROI, F4,1248=4.01, *p*<0.05). There were prominent developmental decreases in MD in face- and character-selective fROIs (Fs>4.13, *p*<0.05), but not in place-selective cortex (F_2,324_=0.39, *p*=n.s.). Additionally, within face-selective fROIs, MD development was observed in deeper cortical layers than superficial layers (F6, 540=4.82, *p*<0.05).

Together, results show that both T_1_ and MD in cortex decrease from age 5 to adulthood suggesting microstructural tissue growth and not tissue loss.

### Development of T_1_ and MD in gray matter correlates with CT

*Does tissue growth in face- and character-selective regions correlate with apparent cortical thinning?* We reasoned that if thinning relates to developments in tissue properties then there will be a significant positive correlation between CT and T_1_ (or MD). To test this, we quantified in each fROI the relationship between CT and T_1_ (or MD) across cortical depths. In face-selective (bilateral pFus-faces and mfus-faces) and character-selective fROIs (bilateral pOTS-chars and left mOTS-chars), we found a significant positive correlation between CT and T_1_ (Rs>0.48, *ps*<0.05) and between CT and MD (Rs>0.36, *ps*<0.05). The highest correlation was between CT and T_1_ at ∼50% cortical depth from the pial surface (**Fig. 4**) and between CT and MD at ∼80% cortical depth (**Fig. S8b**). The correlation between CT with T_1_ remained significant after age was partialled out (**Fig. 4;** same for MD, **Fig. S8b**). In CoS-places, where we did not find development of tissue properties, we also did not find a significant correlation between CT and T1 (right: CoS-places, R=0.3, *p*=n.s.) or CT and MD (Rs<-.06; *p*s=n.s.), except for a significant correlation between CT and T_1_ at ˜50% depth in left CoS-places (R=0.47, *p*<0.05; significant after age partialled out). Together, these analyses suggest that in mid and deep cortical depths of face- and character-selective fROIs, participants with more tissue (lower T_1_ and MD) have a thinner cortex than those with less tissue (higher T1 and MD).

**Figure 4.**
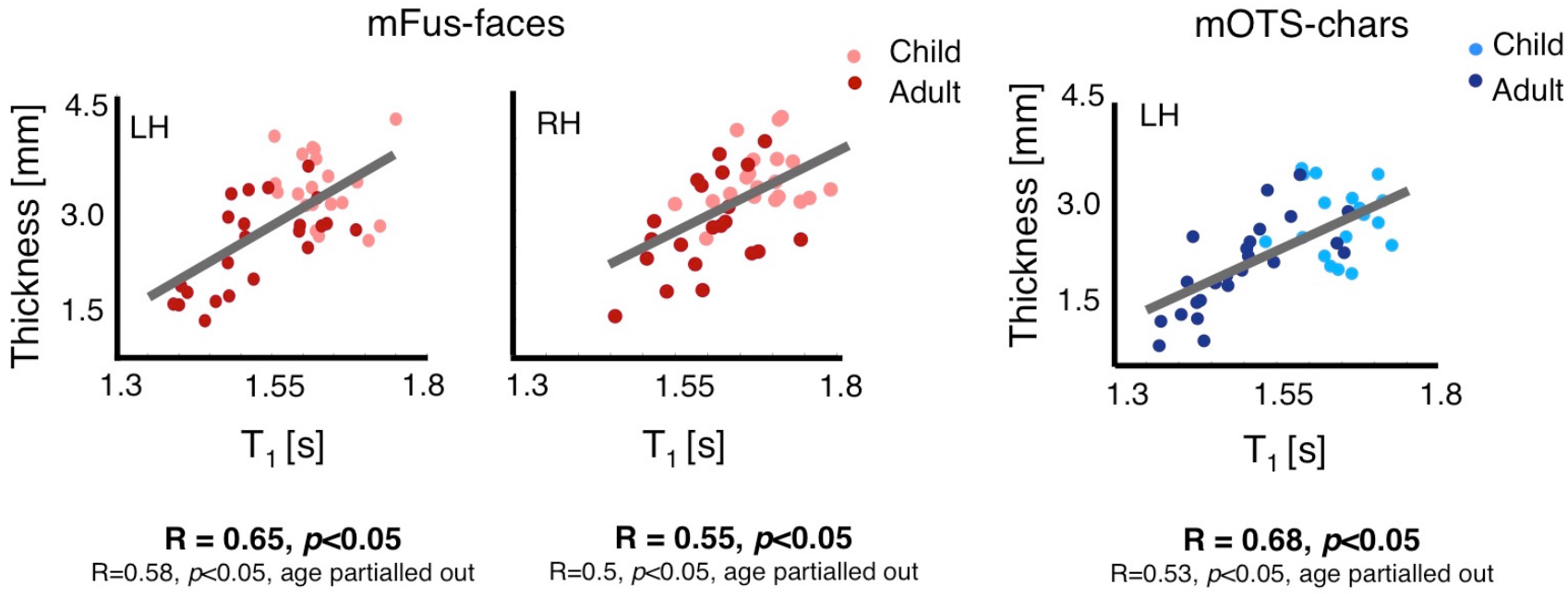
T_1_ and MD in mid and deep cortical depths in face- and character-selective areas correlates with cortical thickness (CT). (a) *Red:* scatter plot showing a significant positive correlation between CT of mFus-faces and T_1_ at mid cortical depth (˜50% projection into gray matter). *Blue:* same for left mOTS- characters. Each point reflects data from one participant. Children are indicated in lighter colors than adults. *LH:* left hemisphere. *RH:* right hemisphere.

### Cortical curvature and surface area are linked to developmental thinning in CoS-places

Our measurements did not find tissue development in CoS-places even as the CoS-places thins from age 5 to adulthood (**Fig. 1b**). Thus, we next asked if cortical thinning in CoS-places is linked to morphological changes including changes in cortical curvature and surface area.

As CoS-places is in a sulcus^33^ and sulcal fundi are typically thinner than sulcal walls and gyri^39^, we examined if developmental changes to the curvature of CoS-places or developmental shifts in the location of CoS-places towards the fundus may explain cortical thinning. In morphological terms, the bottom of the sulcus, or the fundus, has the largest positive curvature (more concavity) than the sulcal wall or a gyrus (less concavity). Analysis of cortical curvature implemented in FreeSurfer revealed that adults’ CoS-places was on a more concave surface than children’s (main effect of age with factors age of subject and hemisphere, F_2,98_=3.47, *p*<0.05, **Fig. 5a**). In contrast, there was no curvature development in face- or character-selective fROIs (Fs<1.45, *p*s=n.s.). Further, we found a significant negative correlation between CT and curvature in CoS-places (**Fig. 5b**, right: R=-0.43, *p*<0.05, significant when age partialled out; left: R=-0.34, *p*<0.05, n.s. when age partialled out). That is, more concave CoS-places was associated with thinner cortex. There was no significant correlation between CT and curvature in bilateral pFus-faces, mFus-faces, and left pOTS-chars (Rs>-0.3, *p*s>.05), but we found a negative correlation in left mOTS-chars and right pOTS-chars (Rs<-0.33, *ps*<0.05; significant when age partialled out). Overall, results suggest that adults’ thinner CoS-places might be situated in the fundus of the CoS as compared to children’s thicker CoS-places.

**Figure 5.**
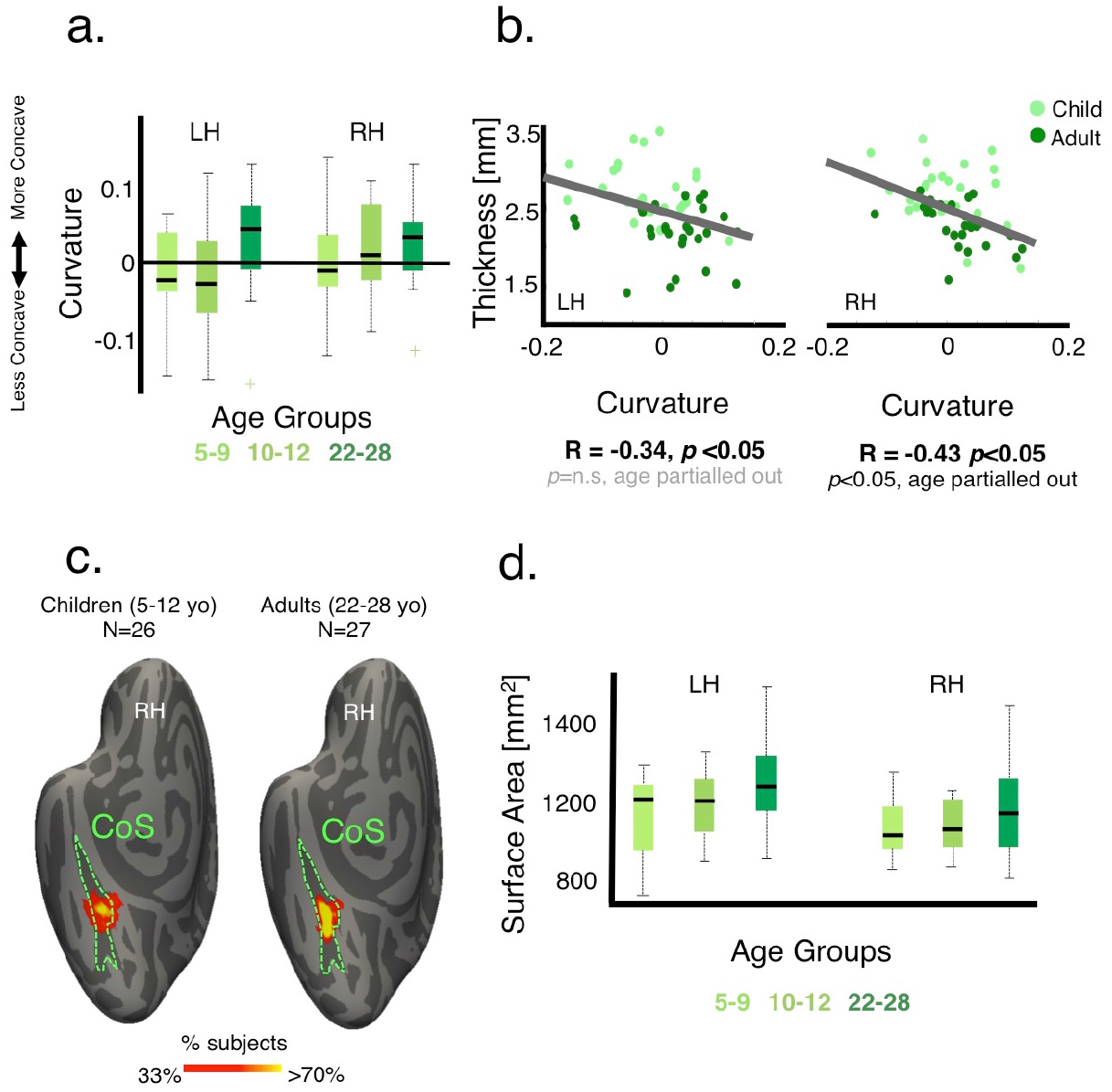
Cortical thickness of CoS-places is linked to development in cortical curvature. (a) Box plots showing median (thick line), 25^th^ and 75^th^ percentile (box), and range (whiskers) of curvature across participants of each age group in bilateral CoS-places. Positive numbers indicate concave surfaces (sulcal folds) and negative numbers indicate convex surfaces (gyral folds). (b) Scatterplot and correlation between CT and cortical curvature of CoS-places. Each dot represents a participant. (c) Probabilistic group maps of the right CoS-places in children (left) and adults (right). Color indicates % participants in which each vertex is included in their CoS-places fROI. Data from both age groups are shown on the FreeSurfer average cortical surface from 39 independent adults^64^. Adults’ CoS-places tends to be localized in the fundus of the CoS. Children’s CoS-places is less consistently found in the fundus and tends to be distributed along the lateral-medial axis of the CoS. (d) Box plots showing median (thick line), 25^th^ and 75^th^ percentile (box), and range (whiskers) of surface area across participants of each age group in bilateral CoS. Surface area of the CoS increases from age 5 to adulthood in both hemispheres. *LH:* left hemisphere. *RH:* right hemisphere. Children are indicated by lighter colors.

To explore this prediction, we visualized the location of CoS-places on the cortical sheet, by generating a group map of CoS-places in children (N=26) and adults (N=27) using cortex-based alignment to the FreeSurfer average brain. As shown in **Fig. 5c** for the right hemisphere, while some of the children’s CoS-places is within the fundus of the CoS, there is variability across subjects along the lateral-medial axis of the CoS. In comparison, adults’ CoS-places is less variable and is consistently localized in the CoS fundus.

Change in structural-functional coupling across development is not associated with growth in the volume of CoS-places (*p*=n.s.). Thus, we tested for morphological changes in the CoS. That is, we examined (i) if cortex stretches during development^42^, and (ii) if areal expansion is coupled with thinner cortex. We derived the CoS from the FreeSurfer anatomical parcellation in each subject, and evaluated if its surface area, including and surrounding CoS-places, develops. Surface area of the CoS expands with age (main effect of age, F_2,100_=4.07, *p*<0.05, **Fig. 5d**) and it was negatively correlated with CT in both hemispheres (Rs<-.41, *ps*<0.05; significant when age partialled out). Together, results suggest that thinning in place-selective cortex is linked to morphological developments whereby the cortex stretches and leads to the location of adult’s CoS-places to be consistently in the fundus.

### Increased myelination as likely mechanism underlying cortical thinning in face- and character-selective areas

*What cellular mechanism underlies tissue growth in face- and character-selective fROIs?* We hypothesized that one tissue compartment that may affect cortical development of T1 and MD may be myelin. However, little is presently known about the myelination of axons in VTC, and how deep into cortex they penetrate. Addressing this question requires measurements of myelin in histological tissue slices of postmortem brains.

While we were unable to obtain pediatric postmortem tissue slices to measure myelin development, we were able to leverage tissue differences across fROIs as a proxy of developmental myelination changes and to examine the effect of myelin on *in vivo* cortical tissue measurement. That is, another aspect of our results revealed that in adults, but not children, relaxation rate (R_1_=1/T_1_) was higher in face-selective fROIs (bilateral pFus-faces and left mFus-faces) than in CoS-places (**Fig. 6a**). In other words, tissue of face- and place-selective regions is undifferentiated in childhood, and development leads to differentiated R_1_ across adult face- and place-selective fROIs. We reasoned that if myelin contributes to the development of R_1_ in face-selective cortex, then face-selective cortex in adults should be more myelinated than place-selective cortex, and especially so in deeper cortical layers.

**Figure 6.**
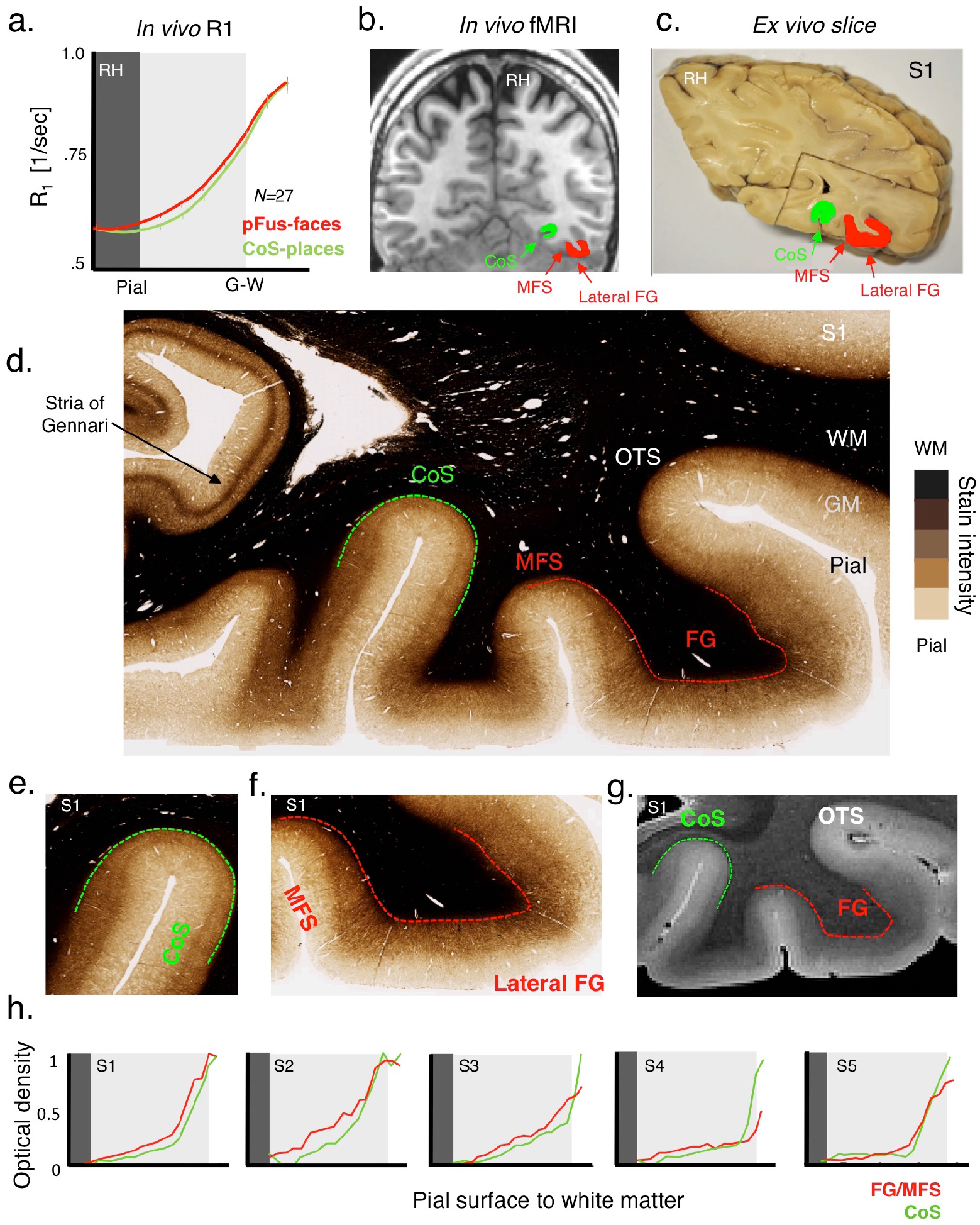
Validation of adult *in vivo* data using adult post-mortem myeloarchitecture. (a) *In vivo* measurement of relaxation rate (R_1_=1/T_1_[1/sec]) shows greater R_1_ in right pFus-faces than CoS-places in middle and deep cortical layers. Data are averaged across 27 adults. *Errorbars*: SEM. (b) *In vivo* example adult coronal slice showing the location of pFus-faces (red) and CoS-places (green) with respect to the mid-fusiform sulcus (MFS), fusiform gyrus (FG), and collateral sulcus (CoS). (c) Photograph of a postmortem section of subject (S1) showing tissue block that was used for histology. *Green*: CoS; *Red:* lateral FG. (d) A sample 30 microns thick histological section of sample S1, stained with modified Gallyas stain for myelin. *Dashed red:* MFS/lateral FG; *Dashed green:* CoS; All slices from each subject in **Fig. S9**. (e-f) Same as (d) zoomed on the CoS (e) and lateral FG (f) to illustrate the penetration of myelin and radial fibers across cortex. (g) High-resolution (50 microns) T2* weighted image of the same section as (d) illustrating differentiation of MR contrast measurements across cortex consistent with the myelin staining in d-f. (h) Measurements of optical density across depths in each of the 5 postmortem brains (S1-S5) for an anatomical section along the CoS (green) and lateral FG (red).

To identify these regions in postmortem histological slices, we need anatomical markers. Prior research from our lab identified reliable anatomical landmarks that predict face-selective and place- selective regions in VTC: the mid-fusiform sulcus (MFS) predicts face-selective regions^32^ and the intersection of the anterior lingual sulcus and CoS predict CoS-places^33^ (**Fig. 6b**). We identified these landmarks in coronal sections (30 microns thick) from 5 postmortem brains that contained the fusiform gyrus (FG), MFS, CoS, and calcarine sulcus (**Fig. 6c**), and then stained histological sections containing these landmarks for myelin using modified Gallyas stain^43^ (**Figs. 6d-f**). We estimated myelin as optical density (OD), which indicates the index of light refracted through the tissue (more myelin produces a darker stain, **Fig. 6d, Fig. S9**).

Myelin stained coronal sections (example: **Fig. 6d;** all brains: **Fig. S9**) revealed striking laminar differences across cortex. Within cortex, deeper cortical layers had higher myelin than superficial layers. The differentiated pattern is also evident in high-resolution (50 microns) T2* weighted image of the same brain (**Fig. 6g**). Zooming in on the CoS (**Fig. 6e**) and FG (**Fig. 6f**), we noted two interesting observations: (1) myelinated fibers project roughly orthogonal to the cortical surface and the depth of myelin penetration varies between FG and CoS. Myelin penetrates about ∼54% into the cortex in FG, but only penetrates ∼36% in CoS. This is consistent with prior findings using equivolume models^44, 45^ that report deeper myelin penetration in cortical layers of gyri than sulci to compensate for thickness. (2) The amount of myelin is greater (darker stains) in FG than in CoS and this difference is not just due to sulcal vs. gyral differences in myelination, as the Stria of Gennari in V1 (**Fig. 6d**) in calcarine sulcus, is strongly myelinated.

We quantified these observations by measuring normalized optical density profiles along equidistance depths from the pial surface to white matter for MFS/FG and CoS in each post mortem brain. Myelin content increased from superficial layers to deeper cortical layers (**Fig. 6h**). Importantly, estimated myelin in MFS/FG was greater than in CoS in depths away from the pial surface in 4 out of 5 brains (**Fig. 6h**), which is largely consistent with our *in vivo* R_1_ data (**Fig. 6a**).

Thus, for the first time, using histological analyses of the MFS/FG and CoS, we show that myelination is a likely mechanism underlying differences in adult *in vivo* T_1_ measurements across these regions. Although we do not have pediatric data, results are consistent with the idea that higher myelination in face- than place-selective cortex in adults is due to increased myelination across development.

## Discussion

Combining CT, qMRI, dMRI, and histology measures for the first time, our study offers the first mechanistic understanding of development of cortical thickness, which is one of the most widely used measures of brain development. First, we found no evidence of pruning after age 5 in any region in VTC. Moreover, in face- and character-selective regions, qMRI and dMRI revealed decreases in T_1_ and MD, both of which occurred in mid and deep gray matter and in the adjacent white matter. These decreases are associated with more tissue, rather than less tissue as predicted by pruning. Second, myelin staining in postmortem brains showed that myelin penetrates into gray matter particularly in the FG, providing evidence that myelin is a significant contributor to T_1_ and MD development in face- and character- selective regions, measured in the present study. Third, we found heterogeneous mechanisms of cortical thinning across VTC. Different than face- and character-selective regions, there was no development of gray or white matter of the CoS-places, instead thinning was coupled with morphological changes. Critically, our results suggest a rethinking of mechanisms of brain development, providing evidence that after age 5, myelin increases in the FG and OTS, and the CoS undergoes morphological changes.

Several innovations in our study have enabled the advancement of understanding mechanisms of cortical thinning across development. First, the use of individual subject analyses with no spatial smoothing and tissue measurements across cortical depths has increased the precision of measurements from centimeters, in prior work employing group analyses, smoothing, and large macroanatomical landmarks^1-5^, to millimeters. Second, leveraging our understanding of functional-structural relationships in VTC^32, 46^, we identified relevant landmarks in postmortem brain slices and validated *in vivo* measurements with ground truth measurements of myelin. Third, we obtained independent measurements of T_1_ from qMRI^18, 19, 47^ and MD from dMRI^21^ in both gray and white matter. These complementary measurements yielded consistent results of growth in both tissue types, strengthening our findings. These advancements not only provide striking empirical evidence supporting developmental theories of increased myelination in deeper gray matter and its adjacent white matter^2,11, 13^, but also underscore the utility of obtaining multiple measurements in the same individual to glean insights into developmental mechanisms.

Our data provide evidence that increased myelination of axons during childhood is a key source of cortical thinning in VTC after age 5. While prior imaging studies documented pervasive thinning throughout childhood, which we replicate (**Figs. 1,S2,S3**), till now the underlying mechanisms have been hypothetical. Some researchers have speculated that cortex thins due to synaptic pruning and cell loss^3^. However, others argued that thinning is likely not due to tissue loss because they found developmental increases in gray and white matter volume^1, 2^. Therefore, they speculated that thinning may be due to a shift in the gray-white boundary from increased myelin. Three of our findings support the latter hypothesis for the development of face- and character-selective regions. First, we found age-related decreases in T_1_ and MD in white matter next to these regions. As myelin explains 90% of the variance in T_1_ of white matter^48^, this finding is likely explained by increased myelination. Second, in cortex, we found decreases in T_1_ and MD far from the pial surface. Third, myelin staining in *ex vivo* adult brains shows that myelin penetrates deep into cortex, especially in the FG. Thus, our findings provide evidence for myelin proliferation into the cortical neuropil during childhood rather than tissue loss, which suggests a rethinking of the term ‘cortical thinning’ to describe structural development of gray matter. In other words, our data suggests that increased myelin during development brightens the gray-white matter interface, shifting the boundary deeper into cortex in adults. While our data suggest that myelin is a key contributor, our prior measurements and simulations indicate that tissue growth affecting T_1_ cannot be exclusively due to myelin increases^26^. Growth in additional microstructures including glia^22, 49^, dendritic arbors, synapses^50^, and iron development in glia and myelin^20^ likely contribute to T_1_ development.

Is it possible that pruning still occurs during development? One possibility is that pruning occurs earlier in development, e.g. during infancy^7, 8^. A second possibility is that pruning after age 5 is smaller in magnitude compared to myelination, and therefore the net effect on MRI measurements is dominated by myelination. Third, it is possible that our voxels and field strength do not have sufficient sensitivity to measure pruning-related developments where the cortex is thin, highly gyrified, or strongly myelinated^20^. Future measurements with high-field MRI and submillimeter resolution, and histology of pediatric and adult postmortem brains can test these possibilities^51^.

Another interesting finding is that different mechanisms underlie apparent thinning in face- and character-selective cortex vs. place-selective cortex, which is merely 2 cm away. In CoS-places, there was no development in either gray or white matter tissue. Instead, CT was correlated with the curvature of CoS-places and the surface area of CoS. This suggests that the CoS may stretch, perhaps deepening the sulcus in adulthood, resulting in cortical thinning. Several mechanistic forces may alter this cortical landscape during childhood, including axonal tension^15^, cytoarchitectural patterning^52^, and differential elasticity properties between white and gray matter^16^. Our findings that morphological changes play a role in cortical thinning, highlight that mechanical forces, an overlooked factor of brain development, should be considered not only during embryonic development, but also during childhood.

What may explain the differential mechanisms of cortical tissue development across VTC? One possibility is that different mechanisms affect development of sulci vs. gyri. However, our data suggest that this is not the case. Both CoS-places and mOTS-chars are in sulci and the latter exhibits tissue growth, while the former does not. Likewise, both CoS-places and V1 are in sulci and the latter has significant myelination (Stria of Gennari in V1, **Fig. 6d**), while the former does not. Another possibility is interplay between functional and structural development. Face-^26,28-30, 53,54^ and character-selective regions^31,54-56^ show a protracted development compared to object- and place-selective regions in VTC. Research on oligodendrocytes and their progenitor cells indicate that development of myelin is activity-dependent^57^. Based on this work and our findings, we propose a new intriguing hypothesis: cortical regions that exhibit protracted functional development will exhibit protracted myelin development, and consequently prolonged apparent cortical thinning. This hypothesis can be tested in future research.

In conclusion, a major goal of neuroscience is to understand mechanisms of brain development. Our study demonstrates the feasibility of evaluating tissue properties in gray and white matter in children and adults using 3T MRI. Additionally, our study is transformative because it underscores the significance of measuring microstructural and morphological changes to brain tissue across the lifespan, rather than just volumetric and thickness measures. Further, as increased myelin during development brightens cortex, our measurements have implications for (i) widespread algorithms estimating CT, brain volume, and tissue type (gray vs. white matter) from MRI, (ii) generating age- appropriate brain atlases, and (iii) large-scale studies of brain development and neuroanatomy including the Adolescent Brain Cognitive Development (ABCD)^58^ study and the Human Connectome Project (HCP)^59^, as these studies will impact future policies that promote the health and well-being of children. Finally, our data have important ramifications for understanding typical^60^ and atypical brain development as well as for clinical conditions implicating myelin and morphology including autism^61^, dyslexia^62^, and multiple sclerosis^63^.

## Authors’ Contributions

VSN designed fMRI experiment, collected and analyzed fMRI, qMRI, dMRI, analyzed post mortem data, and wrote the manuscript; JG contributed to experiment design, MR data collection, and analysis; MB and BJ contributed to the collection and analysis of MR data. ZZ contributed to the analyses of dMRI. SC contributed to defining anatomical ROIs. KSW contributed to discussions on post-mortem data. EK, CJ, and NW contributed to the collection of post mortem brains, MR of post mortem brain, and generation of histological myelin stains. KGS oversaw all components of the study, and wrote the manuscript. All co-authors have read and approved the submitted manuscript.

## Online Methods

### Participants

27 children (14 females, ages 5-12) and 30 adults (11 females, ages 22-28) participated in our study. Children were recruited from the Palo Alto school district, through flyers and online advertisements. All children attended public schools at the time of recruitment. Adult subjects are Stanford University affiliates. MRI data was collected using a 3T scanner in the Center for Cognitive and Neurobiological Imaging (CNI) at Stanford University. All subjects had normal or corrected-to-normal vision and provided written, informed consent. Protocols were approved by the Stanford Internal Review Board on Human Subjects.

### General training and scanning procedures

#### Multiple sessions

Adults and children participated in several scanning sessions on different days, to obtain quantitative MRI (qMRI), diffusion MRI (dMRI), and functional MRI (fMRI) data.

#### Training

Children underwent a separate training session, prior to scanning, inside an MRI scanner simulator. Children were trained to remain still by monitoring their motion, using a motion sensor attached to their forehead and providing them with feedback regarding the amount of head movement they made. Children who stayed still during the mock session were invited back for MRI scans.

#### Acquisition

*In vivo* anatomical and functional data were acquired in each subject using a 3T GE Signa MRI scanner, using a custom-built phase array 32-channel, head coil using methods as in prior studies^18,26^. During MRI scanning, participants lay supine inside the magnet. Visual stimuli were projected onto a monitor and were viewed through an angled mirror mounted above the participant’s head.

### qMRI data

#### Acquisition

Subjects were scanned using spin-echo inversion recovery with an echo-planar imaging (EPI), read-out (SEIR-EPI). This scan was done with a slab-inversion pulse and spatial-spectral fat suppression. We used 2mm^2^ in-plane resolution with a slice thickness of 4 mm and the EPI readout was performed using 2X acceleration. QMRI parameters were measured from spoiled-gradient echo images acquired with different flip angles (α = 4°, 10°, 20° and 30°, TR = 14 ms, TE = 2.4 ms) and a voxel resolution of 0.8×0.8×1 mm^3^, which was resampled to 1 mm^3^ isotropic. For SEIR-EPI, the TR was 3 s. The echo time was set to minimum full; inversion times were 50, 400, 1200 and 2400 ms. Anatomical data were aligned to the AC-PC plane. From the qMRI, we generated whole-brain T_1_-weighted anatomy^18, 26^.

#### Quality assurance

Subjects whose anatomical data appeared blurry or showed ringing due to motion were asked to return for a second scan on a different day (*N subjects* = 4, mean age = 6.75 years). The second scan was of sufficient quality and was used for subsequent visualization and cortical surface reconstruction.

#### General preprocessing and analyses

The spoiled-GE and the SEIR scans were processed using the mrQ software package in MATLAB to produce the T1 and MTV maps. The mrQ analysis pipeline corrects for RF coil bias using SEIR-EPI scans, producing accurate proton density (PD) and T_1_ fits across the brain. MTV maps were produced by calculating the fraction of a voxel that is not MR visible -water (CSF voxels are taken to be nearly 100% water). The full analysis pipeline can be found at (https://github.com/mezera/mrQ)^18^.

#### Reconstruction of cortical surfaces

T_1_ images underwent automated cortical surface reconstruction using FreeSurfer (http://surfer.nmr.mgh.harvard.edu/). The anatomical images were segmented into white and gray matter. White matter surfaces were inspected and manually fixed for missing or mislabeled white matter voxels using ITK-SNAP (http://www.itksnap.org/). A mesh of each participant’s cortical surface was generated from the boundary of the white and gray matter. This mesh was inflated for visualization of activations inside the sulci.

#### Generation of cortical thickness, curvature maps, and surface area

For each subject, we used FreeSurfer’s automated algorithm to define the gray-white matter boundary, pial surface, and cortical thickness (CT) maps. CT (in mm) is measured as the distance between the gray-white boundary and pial surface^39^. FreeSurfer’s algorithm also generates curvature files, which contain information about the amount of curvature at each vertex (i.e., sharper the curve, higher the value) and information whether the vertex is on a sulcus or gyrus (indicated by a sign: a positive sign indicates a sulcus, or concave fold, while a negative sign indicates a gyrus, or convex fold). Mean curvature is made up of the average of principal curvatures derived from the inverse of the radius of the osculating circles at each point on the surface on the gray-white matter junction. Surface area in mm^2^ of the collateral sulcus was calculated from FreeSurfer’s anatomical parcellation of the collateral sulcus for each participant and both hemispheres using FreeSurfer’s *mris_anatomical_stats* and *aparcstats2table* functions.

#### Analysis of CT in FreeSurfer’s anatomical parcellations

To replicate previous results of cortical thinning in large anatomical expanses of the brain^1, 2^, we measured CT in nine anatomical parcellations of the left and right VTC obtained from FreeSurfer’s algorithm for each subject (**see Fig. S3b**). These include: occipital pole, calcarine sulcus, inferior temporal gyrus, occipital temporal sulcus, inferior temporal sulcus and gyrus, collateral sulcus, fusiform gyrus, parahippocampal gyrus, and temporal pole. We generated an “average brain” from our 27 adults and 26 children using FreeSurfer’s *make_average_subject* algorithm. Next, each subject’s brain and their corresponding CT maps were transformed into this average brain space using FreeSurfer’s *mris_preproc* and *mri_surf2surf* algorithms. We then measured average CT per anatomical region, across all subjects in an age group, and conducted a 3-way ANOVA with age of subject, hemisphere, and parcellation as factors.

### dMRI data

#### Acquisition

Consistent with methods in our prior study^41^, in each subject, we obtained two whole-brain, diffusion-weighted, dual-spin echo-sequence (60 slices, TE = 96.8 ms, TR = 8000 ms, 96 diffusion directions, b0 = 2000s/mm^2^, voxel size 2×2×2 mm^3^). Ten non-diffusion-weighted images were collected at the beginning of each scan.

#### Quality assurance

We assessed motion during dMRI from tensor images. Two 5-9 year olds, four 10-12 year olds, and three 22-28 year olds moved during dMRI excessively (motion > 2 voxels, 1 voxel = 2 × 2 × 2 mm^3^). Data from these participants was excluded from subsequent dMRI analyses. MD data is consequently reported for 20 children (ages 5-9, *N*= 9, ages 10-12, *N*=11) and 24 adults (ages 22-28).

#### General preprocessing and analyses

Diffusion data analyses were done using mrDiffusion (http://white.stanford.edu/software). Non-diffusion-weighted images were averaged, and diffusion-weighted (DW)images were registered to this mean image using a two-stage model with a coarse-to-fine approach that maximized mutual information. DWIs were motion and eddy-current corrected and aligned to each subject’s whole brain anatomy using SPM8 (http://www.fil.ion.ucl.ac.uk/spm/software/spm8/). Tensors were fit to each voxel using a least-squares algorithm that removes outliers. For each subject, we obtained mean diffusivity (MD) maps from the tensors files.

### fMRI data

#### Acquisition

Data were acquired using a T2*-sensitive gradient echo spiral pulse sequence with a resolution of 2.4 × 2.4 × 2.4 mm^3^, TR = 1000 ms, TE = 30 ms, flip angle = 76°, and FOV = 192 mm. We collected 48 oblique slices, oriented parallel to the superior temporal sulcus, using a multiplexing technique allowing whole-brain coverage of functional data. The same prescription was used to obtain anatomical T1-weighted images, which were used to align functional data with the whole brain high-resolution anatomical volume of each participant.

#### Functional localizer

In order to localize face-, character-, and place-selective regions of interest (ROIs), subjects participated in a localizer experiment based on our prior methods^26, 37, 53^. Subjects participated in 3 runs of a functional localizer experiment (5.24 min/run) with 78, 4s blocks in each run. Subjects viewed gray-scale stimuli which were blocked by category. Images consisted of two subtypes from each of five categories: characters (numbers and pseudo-words), bodies (limbs and headless bodies), human faces (child faces and adult faces), places (houses and indoor scenes), and objects (guitars and cars) (see **Fig. S1a** for example images per category). Each image was shown only once during the experiment. In each 4 s block, different stimuli from one of the above categories were shown at a rate of 2 images per second. Blocks were counterbalanced across categories and also with baseline blocks consisting of a blank, gray background. During the scan, subjects fixated on a central dot and pressed a button when a phase-scrambled, “oddball” image appeared randomly in a block (˜33% of the blocks).

#### Quality assurance

(1) Motion. Data of each subject were corrected for within-run and between-run motion. Only runs with motion of less than two voxels were included in the study (as in our prior studies^26,53^). To test for development, we performed a 2-way ANOVA with factors, age of subject and motion type. (2) Time series signal to noise ratio (tSNR). As a control to test if developmental changes were driven by smaller signal to noise ratio in children than adults, we measured tSNR across children’s and adults’ left and right VTC ROIs. For each subject, we defined left/right VTC as the boundary between the posterior transverse collateral sulcus (ptCoS) to the anterior tip of the MFS, including the extreme edges of the CoS and OTS. tSNR was computed for each voxel as follows: tSNR = mean(time series)/standard deviation (time series) and then averaged across all voxels of the VTC, across runs and across subjects in an age group. We performed a 2-way ANOVA with factors, age of subject and hemisphere to test development in tSNR.

#### General processing and analyses

Localizer data were analyzed using code written in a MATLAB based mrVista toolbox (http://github.com/vistalab) as in our prior publications^26, 37, 53, 54^. Data were not spatially smoothed and were analyzed in each subject’s native brain space. The time courses of each voxel were converted from arbitrary scanner units into units of % signal change.

#### General Linear Model (GLM)

For each subject’s data, we ran a GLM to model each voxel’s time course. The experimental design matrix was convolved with the SPM hemodynamic response function (HRF) (http://www.fil.ion.ucl.ac.uk/spm) to generate predictors. Using a GLM to fit the predictors to the data, we estimated the response amplitudes for each condition (betas) and residual variance of each voxel’s time course. We used beta values and residual variance from the GLM to generate contrast maps comparing responses in different conditions.

#### Definition of functional regions of interest (fROIs)

Functionally selective voxels were defined as voxels that responded more to images of a selected category than to images of other categories (t>3, voxel level) during the localizer scan. Thus, in each subject, we defined the following regions in the ventral stream. Spatially contiguous clusters of face-selective voxels that responded more strongly to faces (child faces and adult faces) than other stimuli and were located in the posterior lateral fusiform gyrus were defined as pFus-faces. Face-selective voxels near or overlapping the anterior tip of the mid-fusiform sulcus (MFS) were defined as mFus-faces (pFus- and mFus-faces are collectively referred to as the fusiform face area, FFA^34^). Clusters of voxels that responded more strongly to places (house and corridors) than other stimuli and were located in the collateral sulcus (CoS) were defined as place-selective CoS-places also referred to as the parahippocampal place area (PPA)^38^. Clusters of voxels that responded more strongly to characters (numbers and pseudo-words) than other stimuli and were located in the posterior OTS were defined as pOTS-characters, also referred to as visual word form area 1{Stigliani, 2015). More anterior character-selective voxels in the occipital temporal sulcus were defined as mOTS-characters also referred to as visual word form area 2^37^ (mOTS- and pOTS-words are collectively referred to as the visual word form area, VWFA^36^).

### Behavior

We collected behavioral data during the scan in an odd-ball task, in which participants were asked to press a button when they viewed a scrambled image on the screen, which appeared in 33% of the trials. Due to equipment malfunction, however, we were able to collect data from 10 children (ages 5-12, 2 females) and 17 adults (ages 22-26, 8 females). Performance accuracy in the oddball task during fMRI was not significantly different (t_(24)_=0.42, *p*=n.s., two-tailed) across children (78.0±7.78%) and adults (81.43±4.51%).

### In vivo analyses examining anatomical features of functional regions

#### Analysis of CT in functional ROIs of VTC

Data were analyzed using a MATLAB-based, mrVista toolbox. In brief, each subject’s CT surface maps from FreeSurfer’s algorithm were transformed to volume nifti files using FreeSurfer’s *mri_surf2vol* algorithm. For each subject, we averaged CT per fROI and hemisphere, across all subjects within an age group. We performed a 3-way ANOVA with factors, age of subject, fROIs, and hemisphere.

#### Analysis of T_1_, MTV, and MD in functionally-defined white matter (FDWM)

Each individual subject’s fROI was dilated into its neighboring white and gray matter by 5mm, to obtain T_1_, MTV, and MD measures of its functionally-defined white matter. Average T_1_, MTV, and MD were calculated for these FDWM voxels, and averaged across subjects within an age group. We performed a 3-way ANOVA with age of subject, hemisphere, and fROIs as factors per measure.

#### Analysis of T1 /MD by intra-cortical depth

Using FreeSurfer’s *mri_surf2vol* algorithm, each subject’s FreeSurfer-based white matter surface was converted to volume niftis. The volume was projected into gray matter at increasing 20% steps, starting from two locations below the gray-white boundary (to include some white matter just below the boundary) to two locations above the pial surface (to include the entire pial surface). As thickness of the cortical ribbon varies between 2mm to 4mm in the VTC, each increasing depth is between 0.4 mm – 0.8 mm from the gray-white surface normal. For each subject’s dilated fROI, we averaged T_1_ values of those voxels that intersected with each cortical layer, and obtained T_1_ values per cortical depth. We performed 4-way ANOVAs with age of subject, fROI, hemisphere, and cortical depth as factors. Similar analysis was conducted for MD, with the exception that the cortex was divided into 8 steps rather than 10 steps as MD maps were 2mm voxel resolution. For the statistical analysis of variance using depth as a factor, we used every other step instead of all steps to ensure that they reflect data from non-overlapping voxels (5 out of 10 steps for T_1_ and 4 out of 8 steps for MD). For each fROI, we correlated each subjects’ CT with T_1_ (or MD) at each cortical depth (Pearson’s correlation coefficient).

#### Correlation between anatomical curvature and CT

Each subject’s curvature maps from FreeSurfer’s autosegmentation algorithm were transformed into volume nifti files using the FreeSurfer-based *mri_surf2vol* algorithm. Data were analyzed using MATLAB-based mrVista toolbox. Per participant, we calculated average curvature in each fROI and examined if there was a significant correlation (using Pearson’s correlation coefficient) between CT and curvature per fROI.

### Post mortem tissue data

#### Histology (post mortem tissue blocks)

Blocks of five post mortem human brains were provided by the Brain Banking Centre Leipzig of the German Brain-Net (GZ 01GI9999-01GI0299), operated by the Paul Flechsig Institute of Brain Research (University of Leipzig). Information on sex, age of death, cause of death and post mortem interval before fixation (PMI) is provided in **Supplementary Table 2**. The entire procedure of 1) case recruitment, 2) acquisition of the patient’s personal data, including protocols and informed consent forms, 4) the performance of the autopsy, and 5) the handling of the autopsy material have been approved by the responsible authorities. Following the standard Brain Bank procedures, tissue blocks from occipital cortex were extracted, immersion-fixed in 4% paraformaldehyde in phosphate buffered saline (PBS) pH 7.4 for at least 6 weeks. Prior to MR scanning, blocks were washed in PBS with 0.1% sodium azide for at least 24 h in order to avoid biasing the effects of paraformaldehyde on MR relaxation rate. All tissue blocks contained a part of the calcarine fissure, collateral sulcus, and fusiform gyrus.

#### MR Scanning

MR scanning of post mortem tissue samples was performed on a 7T Siemens scanner (Magnetom, Siemens, Erlangen, Germany) equipped with a custom made 2-channel transmit-receive coil with an inner diameter of 6 cm. Samples were placed in a spherical container filled with Fomblin (Sigma Aldrich).

#### Tissue sectioning

For histological processing, tissue blocks were cryoprotected in 30% sucrose/PBS with 0.1% sodium azide. Sections of 30 microns thickness were made on a cryomicrotome (Leica SM2000R with freezing unit Zeiss KS34) and collected in PBS with 0.1% sodium azide.

#### Semi-quantitative myelin mapping in histological sections

For myelin staining, every 20^th^ section was taken out of the series (in total: 6 to 9 sections per case), mounted on glass slides, and air dried. Sections were rehydrated in distilled water and stained following the classical protocol of the Gallyas method^43^ with slight adaptions^65^. In order to assure the complete visualisation of all myelin fibers, including those that are very thin, the developing step was extended to 40 min. After the procedure, slides were thoroughly washed in distilled water, dehydrated and cover slipped with Entellan/Toluene (Merck). Imaging was performed with a Zeiss Axio Scan, Z1 slide scanner with a 10x objective. All images were obtained under equal conditions to ensure comparable data for the optometric processing. Obtained RGB maps were logarithm transformed using the Lambert-Beer transformation and averaged across three color channels with corresponding weights to calculate optical density (OD) of the sections (i.e., OD =0.8*R + 1.2*G+ 1.5*B, where R, G, B are the red, green, and blue channels in a .tiff image file). Thus, OD is proportional to the density of staining chromophore and therefore, reflects local myelin density in the tissue.

### Adult histology data vs. adult in-vivo T_1_ data in face- and place-selective cortex

#### General processing and analyses

Our strategy was to compare adult histology data to adult *in vivo* T_1_ data in face- and place-selective cortex. Thus, for each individual stained slice, we divided the cortex into equidistant depths from the pial surface to white matter. We achieved this by drawing a mid-line in MFS/FG and CoS (along its thickness) and projecting equidistant depths along the mid-line normals in two steps: 1) from mid-line to pial-surface and 2) mid-line to white matter. We then evaluated average OD at each depth (OD at each point was normalized with respect to OD in white matter), which in turn, reflects the measure of the amount of myelin per region.

#### Data availability

All code relevant to data analysis as well as the source data for the main findings (**Figs. 1-6**) will be made available upon request. Majority of the code used in this study was derived from functions available through the open-source code library: https://github.com/vistalab/vistasoft.

## Supplementary Materials

**Supplementary Table 1.** Number of face-, character-, and place-selective functional regions of interest localized in left and right hemispheres of children (ages: 5-9 years, *N*=11, and ages: 10-12 years, *N*=15) and adults (ages: 22-28 years, *N*=27).

| LEFT HEMISPHERE | 5-9<br>years | 10-12<br>years | 22-28<br>years | RIGHT<br>HEMISPHERE | 5-9<br>years | 10-12<br>years | 22-28<br>years |
| --- | --- | --- | --- | --- | --- | --- | --- |
| pFus-faces/FFA1 | 9/11 | 13/15 | 20/27 | pFus-faces/FFA1 | 7/11 | 11/15 | 23/27 |
| mFus-faces/FFA2 | 8/11 | 12/15 | 23/27 | mFus-faces/FFA2 | 7/11 | 14/15 | 21/27 |
| pOTS-chars/VWFA1 | 8/11 | 12/15 | 27/27 | pOTS-chars/VWFA1 | 5/11 | 11/15 | 25/27 |
| mOTS-chars/VWFA2 | 7/11 | 11/15 | 24/27 | mOTS-chars/VWFA2 | 0/11 | 4/15 | 10/27 |
| CoS-places/PPA | 10/11 | 15/15 | 27/27 | CoS-places/PPA | 10/11 | 15/15 | 27/27 |

**Supplementary Table 2.** Information on five post mortem tissue samples used in this study including hemisphere, age, gender, cause of death, and post mortem interval before fixation (PMI).

| Sr. No. | Hemisphere | Age | Gender | Cause of death | PMI |
| --- | --- | --- | --- | --- | --- |
| 1. | Right | 68 | M | Rectal carcinoma | 24 |
| 2. | Right | 79 | M | Heart failure | 48 |
| 3. | Right | 57 | F | Thrombosis | 24 |
| 4. | Right | 74 | F | Heart failure | 24 |
| 5. | Right | 57 | M | Liver carcinoma | 24 |

**Supplementary Figure 1.**
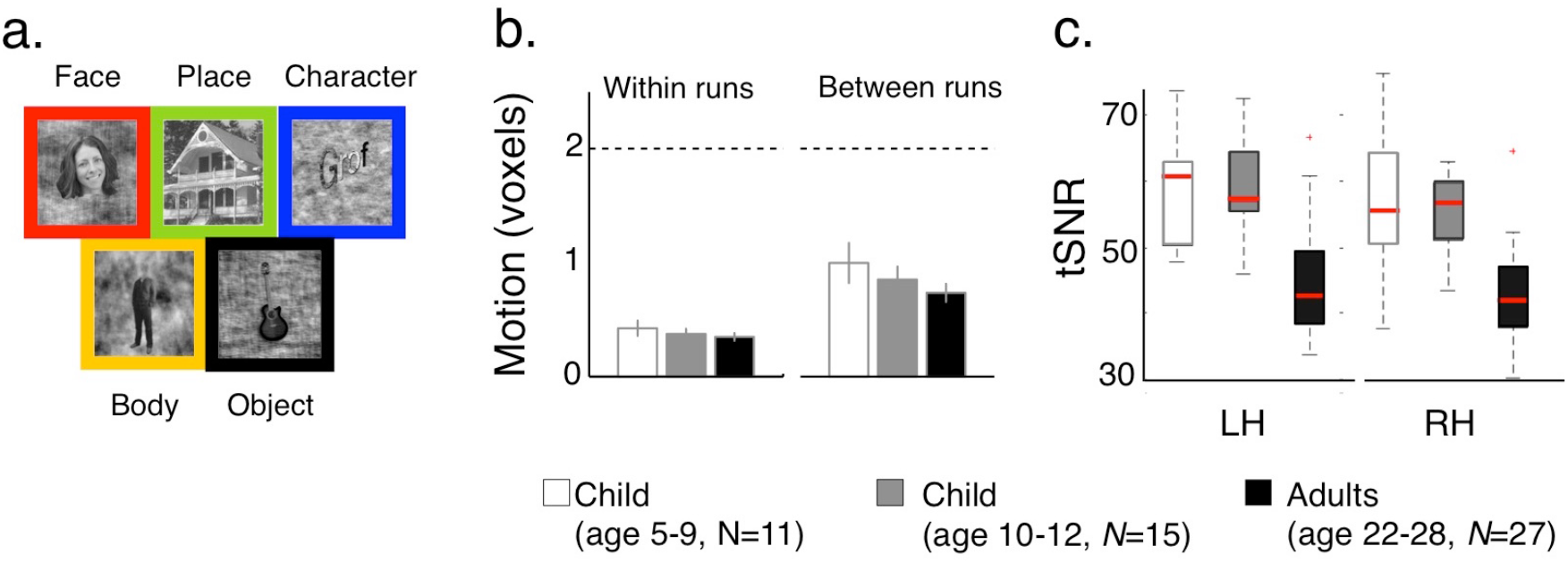
Stimuli used in the functional localizer experiment and experimental controls. (a) Sample face, place, object, body, and character stimuli used in the localizer experiment to identify category-selective regions in children and adults, (b) Graph shows within-run and between-run head motion across three age groups (ages: 5-9, 10-12, 22-28 years). Head motion is comparable across subjects (F_2,100_=2.61, *p*=n.s.). *Error bars:* standard error of the mean (SEM) across subjects within an age-group, (c) Box plots showing time series signal-to-noise-ratio (tSNR) across VTC voxels. *Red line*: median; *box edges:* 25^th^ and 75^th^ percentile, *whiskers:* range; *red crosshairs:* outliers. Children’s tSNR were significantly larger than adults’ across hemispheres (main effect of age, *F*_2,100_=37.49, *p*<0.05). *LH:* left hemisphere. *RH:* right hemisphere.

**Supplementary Figure 2.**
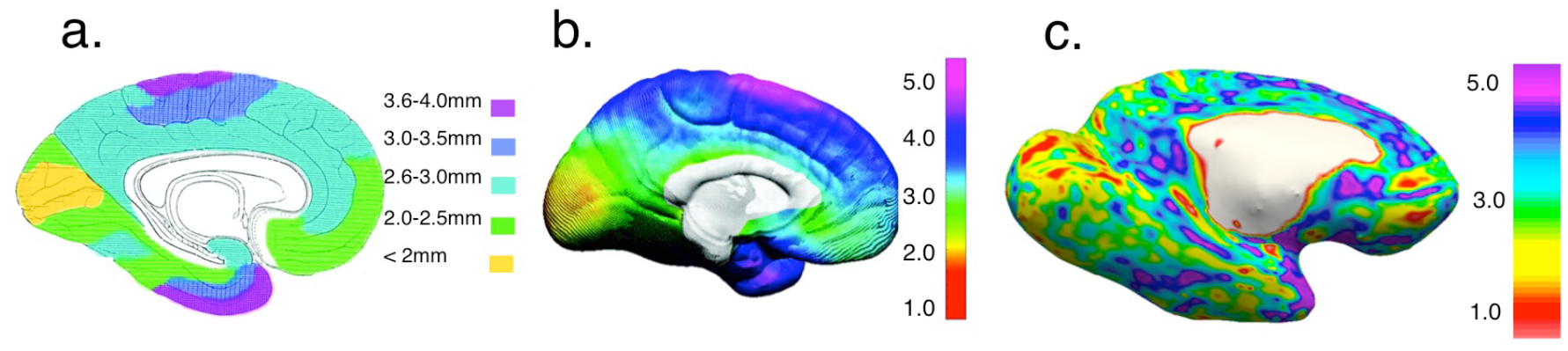
Consistent estimates of cortical thickness (CT) across an *ex vivo study, in vivo* study, and our data. (a) CT map from an adult *ex vivo* data, adapted from the Von Economo (1929). (b) Average CT map from *in vivo* data of 45 adults obtained on a 1.5T MRI scanner, adapted from Sowell et al., (2004). (c) CT map from our *in vivo* data in an example adult scanned in a 3T MRI scanner. Similar color coding is applied across the three CT maps to allow comparisons across data sets. Data are shown on a medial view of the left hemisphere. In general, studies show consistent CT. Medial regions in the occipital lobes are the thinnest, and anterior temporal cortex and superior parietal cortex are the thickest.

**Supplementary Figure 3.**
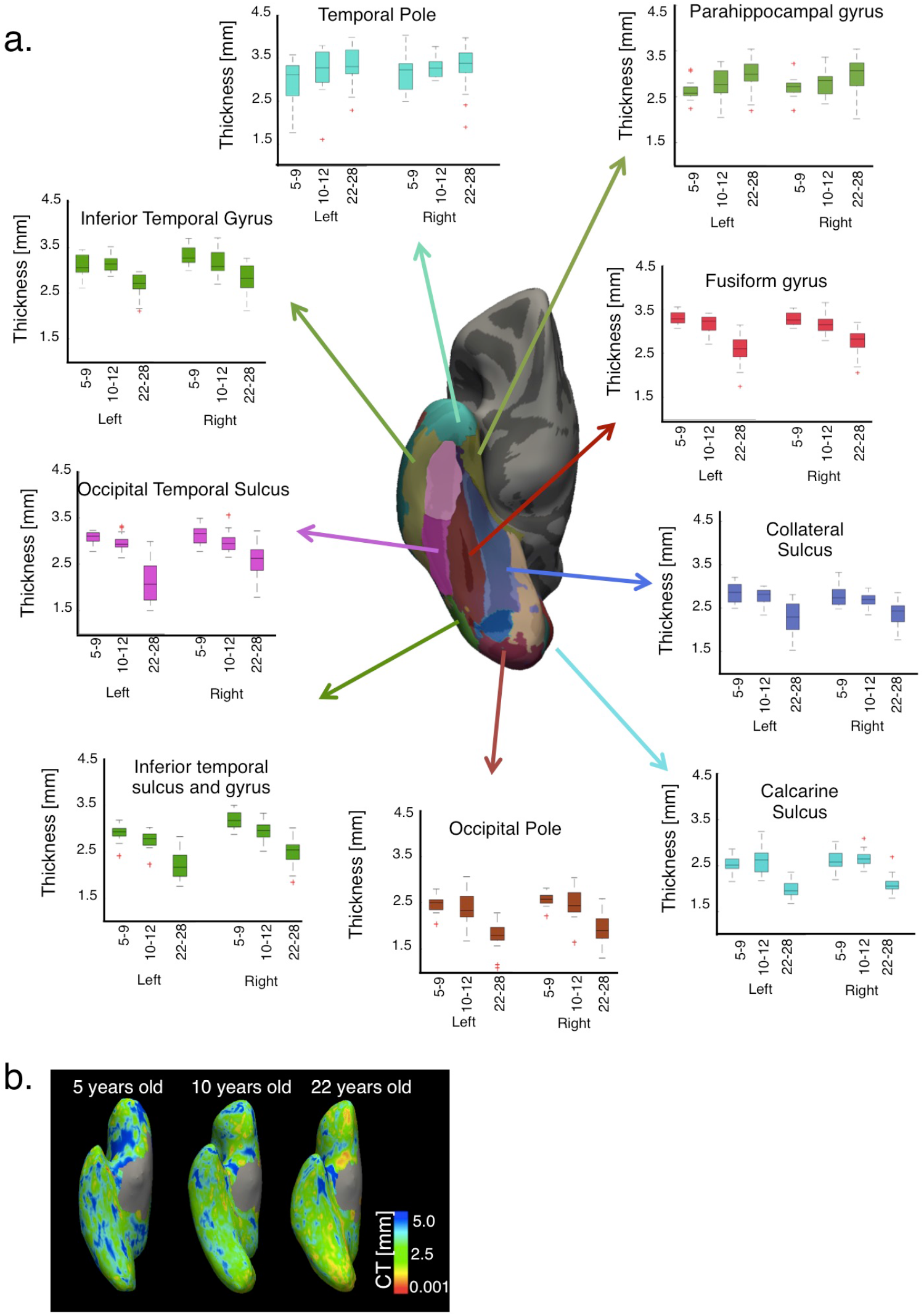
Cortical thinning in anatomical parcellations of the VTC. (a) Development of CT across 3 age groups (ages 5-9, 10-12, 22-28 years) in nine broad anatomical parcellations of ventral occipital and temporal cortex defined in Freesurfer. The 9 parcellations are shown in different colors on an average cortical surface generated across the 53 subjects of our study. Boxplots show median, 25^th^ and 75^th^ percentile, and range of CT in each age group. *Red crosshairs*: outliers. CT significantly decreased from age 5 to adulthood in 7 out of the 9 anatomical parcellations (significant interaction between age of subject and anatomical parcel, *F*_16,900_ = 19.48, *p* < 0.05), with the exception of the temporal pole and parahippocampal gyrus, which showed increased CT with age. (b) CT maps calculated in Freesurfer in three example participants (ages 5, 10, and 22 years) showing CT in millimeters [mm].

**Supplementary Figure 4.**
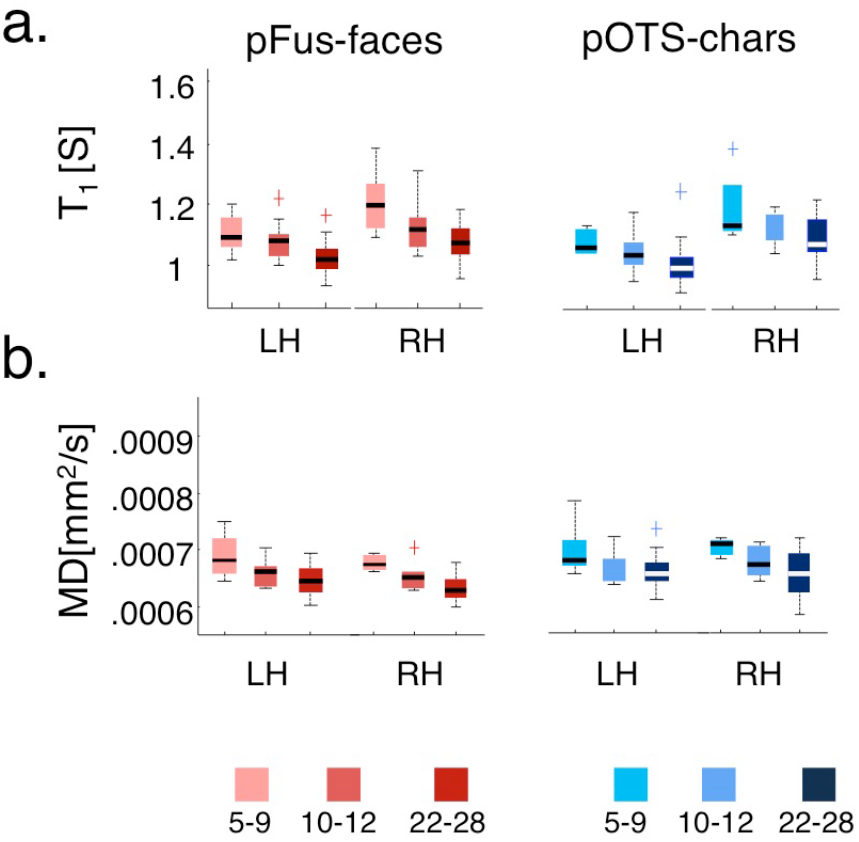
T_1_ and MD in white matter adjacent to pFus-faces and pOTS-characters decrease with age. (a-b) Box plots showing T_1_ and MD in a face-selective regions in the posterior fusiform gyrus (pFus-faces, red) and a character selective region in the posterior occipito-temporal sulcus (pOTS-characters, blue) in children and adults. *Light colors*: 5-9 year olds; *medium colors:* 10-12 year olds; *dark colors:* 22-28 year olds. *LH:* left hemisphere. *RH:* right hemisphere.

**Supplementary Figure 5.**
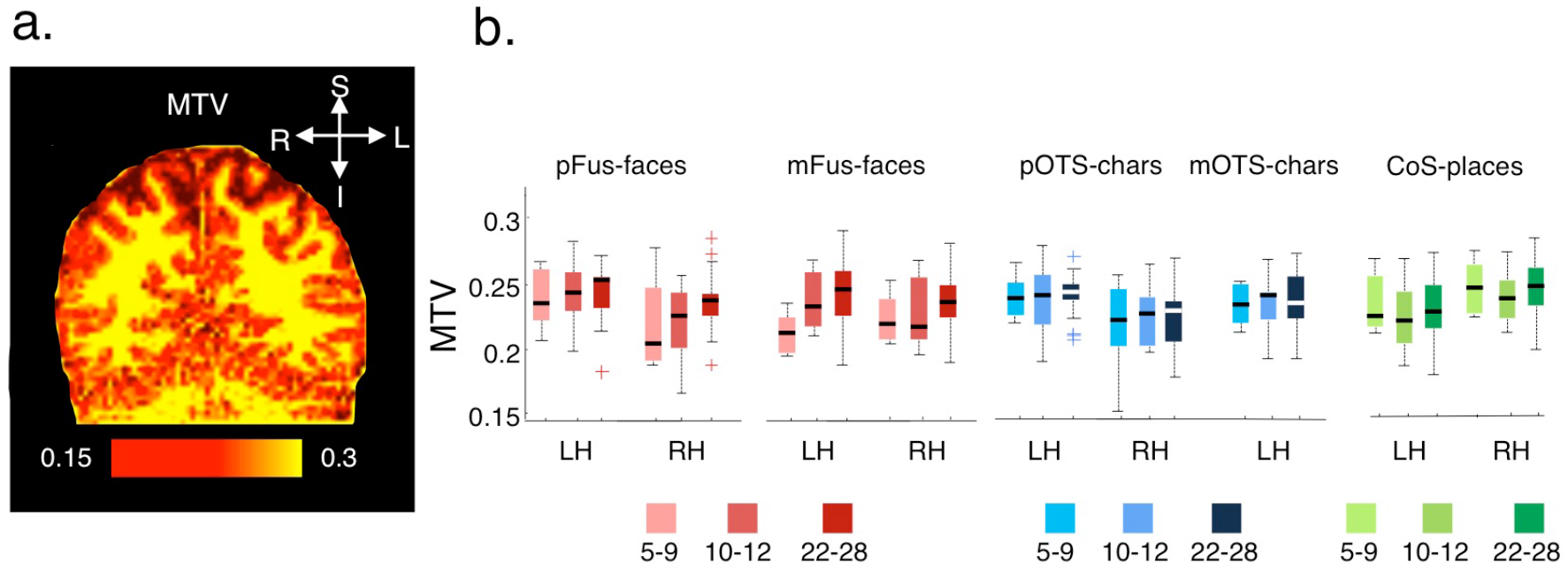
Developmental increase in macromolecular tissue volume (MTV). Coronal slice in an example adult brain showing MTV map. (b) Box plots showing MTV in faceted), character- (blue), and place-selective (green) regions. Boxplots show median (red), 25^th^ and 75^th^ percentile (box edges), range (whiskers) and outliers (red crosshairs) of CT in each age group. MTV progressively increased from 5-9 year olds, to 10-12 year olds, to adults in VTC fROIs (main effect of age: F_2, 384_ = 3.36, *p* < 0.05). *Light colors*: 5-9 year olds, *medium colors:* 10-12 year olds; *dark colors:* 22-28 year olds. *LH:* left hemisphere. *RH:* right hemisphere.

**Supplementary Figure 6.**
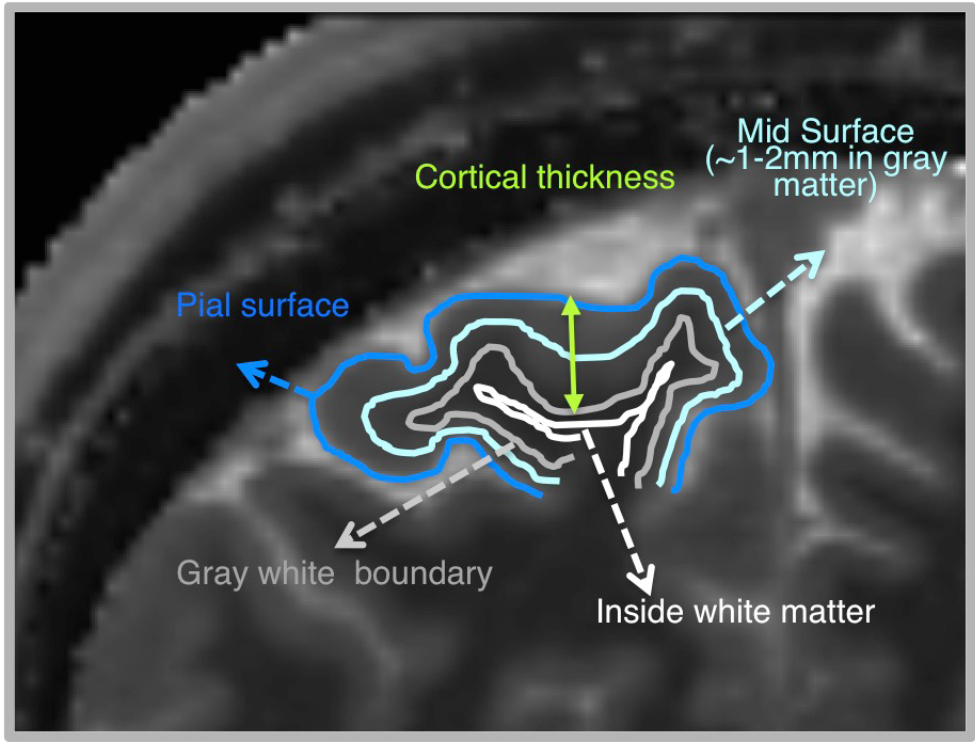
Schematic showing sample projections along the cortical depth. Schematic shows sample projections along the cortical depth used for measuring T_1_ and MD: starting from the pial surface (dark blue line), midsurface (∼50% depth from the pial surface, light blue line), gray-white boundary (gray line) and inside the white matter (white line). The distance between the pial and gray white surfaces is defined as cortical thickness in mm.

**Supplementary Figure 7.**
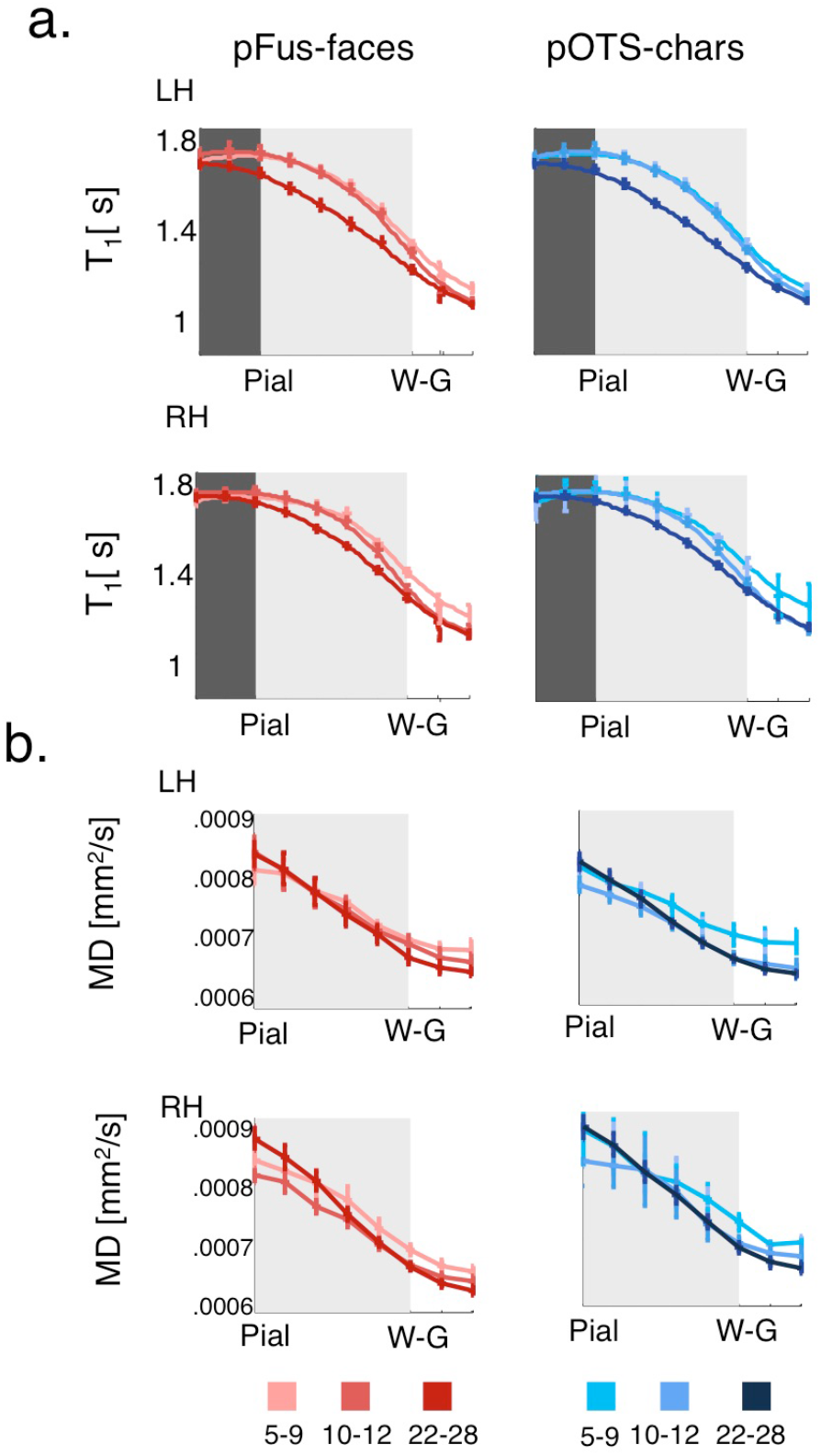
Development of T_1_ and MD of pFus-faces and pOTS- characters as a function of intra-cortical depth. (a) T_1_ curves across equidistance intracortical depths from pial surface to local white matter tissue in bilateral face-selective pFus-faces (red) and character-selective regions pOTS-characters (blue) in three age groups. Development is observed in both face- and character-selective fROIs away from superficial pial surface, in middle cortical depths, (b) MD curves across equidistance intracortical depths from the pial surface to white matter in bilateral pFus-faces and pOTS- characters. Like T_1_, development in MD is observed in both face- and character-selective fROIs. In all plots, *light colors:* 5-9 year olds; *medium colors:* 10-12 year olds; *dark colors:* 22-28 year olds. *Error bars:* SEM. *LH:* left hemisphere. *RH:* right hemisphere.

**Supplementary Figure 8.**
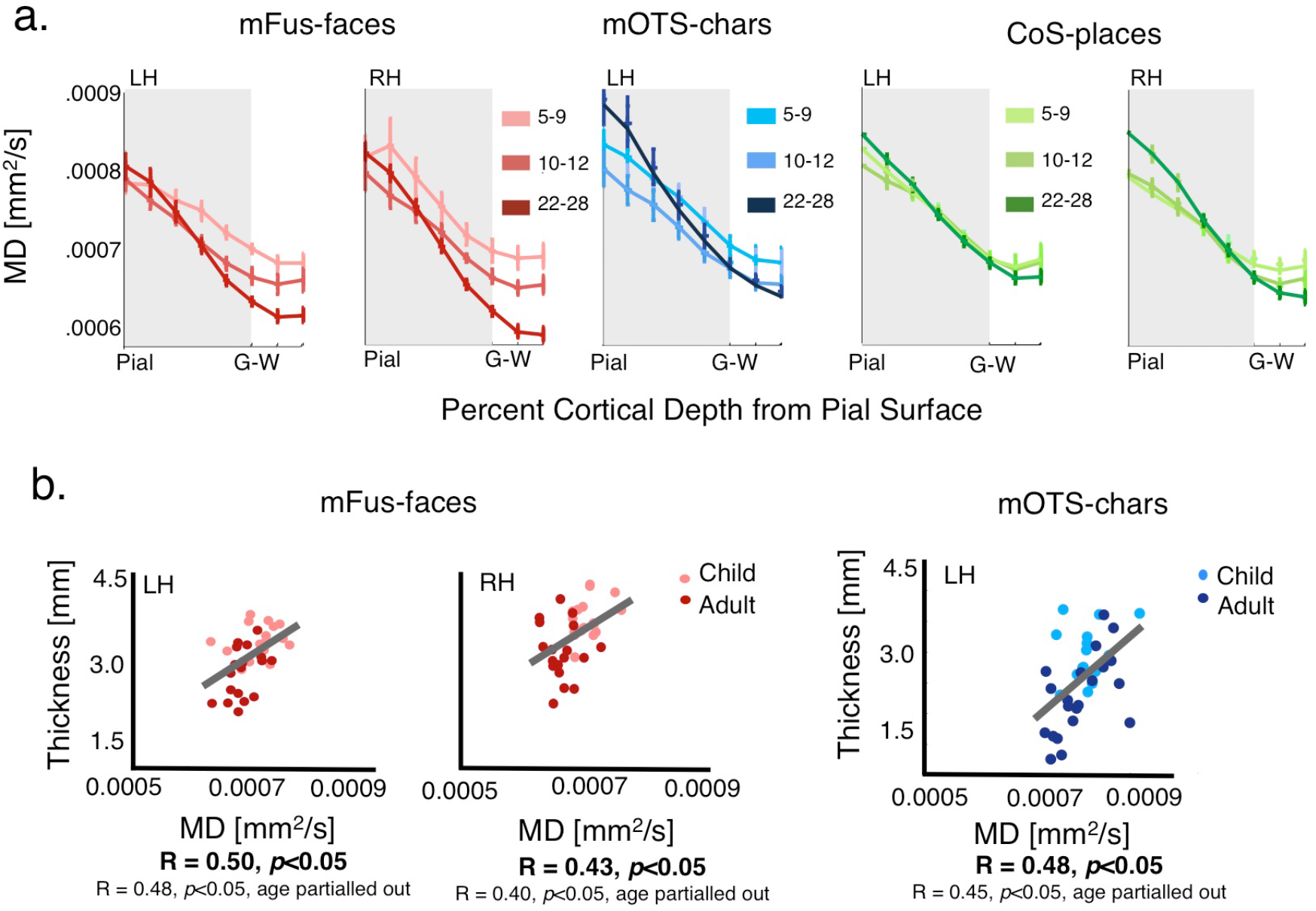
Development of cortical MD as a function of cortical depth and its relationship with cortical thickness. (a) MD curves across equidistance intracortical depths from pial surface into white matter in bilateral face-selective mFus-faces (red), left character-selective regions mOTS-characters (blue), and bilateral place-selective CoS-places (green) in the three age groups. Development is observed in face- and character-selective fROIs, but similar developmental pattern was not found in place-selective fROIs (a significant interaction between age of subject and fROI, F_4, 2496_%6.81, *p*<0.05). In all plots, *light colors:* 5-9 year olds; *medium colors:* 10-12 year olds; *dark colors:* 22-28 year olds. *Error bars:* SEM. *LH:* left hemisphere. *RH:* right hemisphere, (b) *Left:* Positive correlation between CT and MD of mFus-faces; *Right:* Positive correlation between CT and MD of left mOTS-chars. MD is measured at ∼80% depth. Results indicate that tissue growth in deeper cortical layers of face- and character-selective fROIs is linked to their cortical thinning. No significant relationship was found between CT and MD in place-selective cortex. Each dot is a participant; *light color:* children; *dark colors:* adults.

**Supplementary Figure 9.**
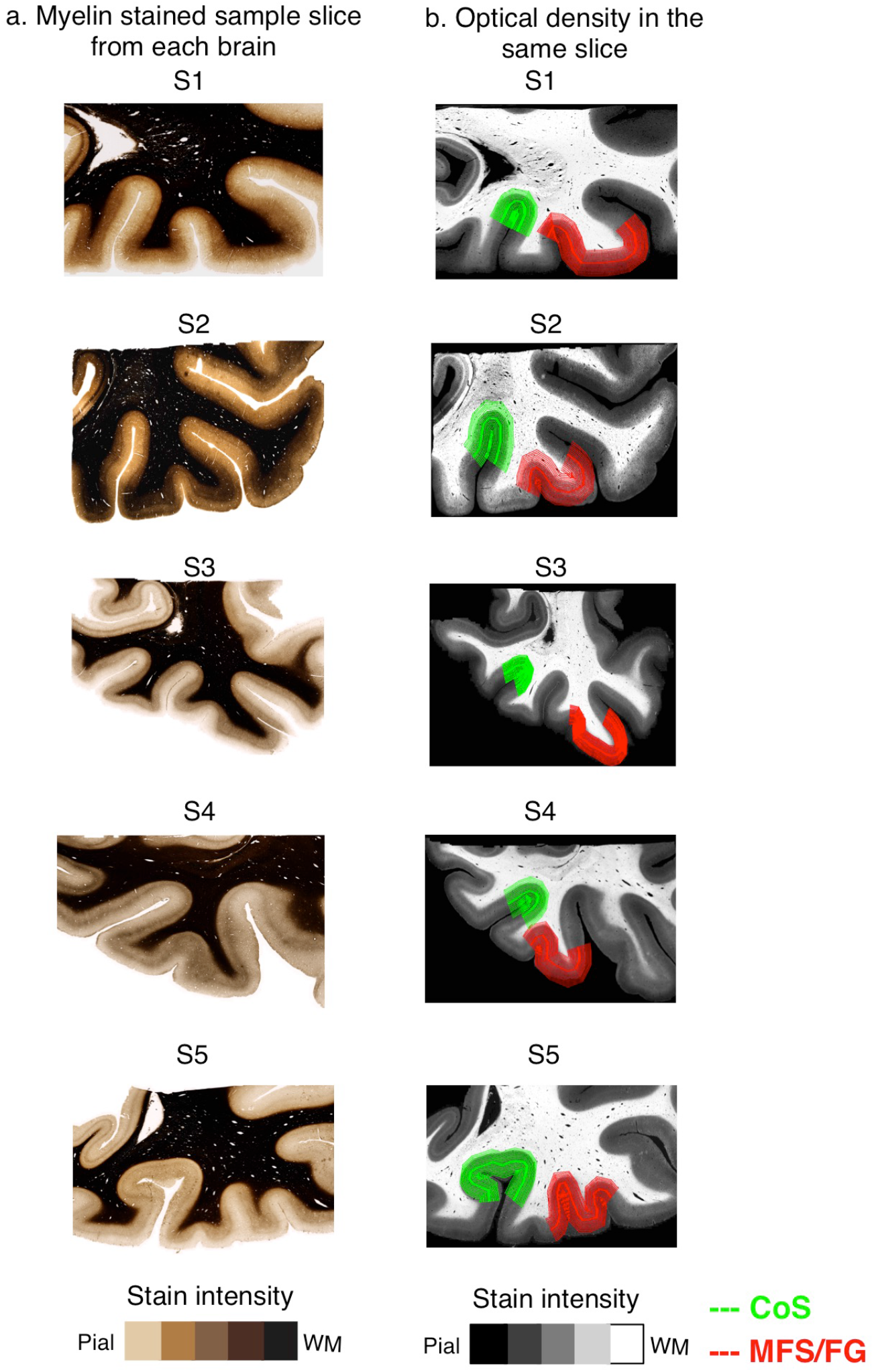
Myelination in histological slices containing the collateral suclus (CoS) and fusiform gyrus (FG). (a) Sample 30-micron thick histological sections in each of the 5 postmortem brains (S1-S5) stained with modified Galiyas stain for myelin. Darker colors indicate higher myelin content, (b) Optical density measurements in the same slices as in (a) showing the center-depth cortical path along which measurements for CoS (green) and MFS/FG (red) were extracted.

## Acknowledgements

This research was funded by NIH grants 1RO1EY 02231801A1 and 1R01EY02391501A1 to KGS, 5T32EY020485 to VSN, and NRSA F31EY027201 to JG. We thank Prof. Dr. Thomas Arendt from the Paul Flechsig Institute of Brain Research, University of Leipzig, for providing post mortem brain tissue samples. The project was supported by the European Research Council under the European Union’s Seventh Framework Programme (FP7/2007-2013) / ERC grant agreement n° 616905, and the BMBF (01EW1711A & B) in the framework of ERA-NET NEURON.

